# Temperature and flow impacts on spawner-to-fry production in Okanagan sockeye salmon

**DOI:** 10.64898/2026.09.25.754467

**Authors:** Patrick L. Thompson, Karilyn Alex, Howard W. Stiff, Athena D. Ogden

## Abstract

Okanagan sockeye salmon have shown a strong recovery over the past three decades, but the contribution of environmental conditions and management actions to this recovery has not been determined. We developed a hierarchical Bayesian model linking spawner abundances to fry emergence observations to evaluate how flow and water temperature during adult upstream migration and incubation periods influence spawner-to-fry production. Production declined by 62% in years with earlier onset and longer duration of periods when warm river temperatures created barriers to adult migration, indicating that migration conditions can have carry-over effects on reproductive success even when spawners successfully reach the spawning grounds. Additionally, production was 46% lower when flows fell above or below the management target range designed to minimize scour and desiccation during egg incubation. In contrast, we found little support for effects of pre-spawning flows or incubation temperature on production. These findings provide quantitative evidence that current flow management is increasing freshwater spawner-to-fry production and has likely contributed to the Okanagan sockeye recovery. Effective flow management may therefore help offset climate-driven impacts that increasingly threaten this population.

## Introduction

Sockeye salmon (*Oncorhynchus nerka*) populations are in decline over much of their southern range in North America (Rand et al. 2012; Grant et al. 2019), highlighting the urgent need for management measures that can boost survival and contribute to maintenance and recovery. In this context, the Okanagan sockeye represent a rare success story where targeted management measures have proven effective (Alexander et al. 2024), resulting in a twentyfold increase in spawner abundance since the early 2000s (Bailey et al. 2025). However, because multiple management measures were implemented concurrently—including habitat restoration, hatchery restocking, improved dam passage, and management of dam outflows to ensure “fish-friendly flows”—alongside changing environmental conditions, it has proven challenging to quantify the contribution of individual interventions. Based on a weight-of-evidence approach, Alexander et al. (2024) concluded that these management interventions jointly contributed to the recovery but that a shift towards more favourable coastal marine conditions in 2007 also played a role. Still, quantitative estimates of how environmental conditions and individual management interventions impact production and survival are needed in order to ensure that limited conservation resources are used effectively, and to assess whether similar interventions could support recovery in other salmon populations.

This study focuses on the spawner-to-fry component of the Okanagan sockeye life cycle, with the goal of quantifying how environmental conditions and management interventions influence production at this stage. Specifically, it focuses on the Osoyoos Lake population, which was the only remaining sockeye population in the Okanagan prior to the recent reintroduction into Skaha and Okanagan Lakes and which remains the most abundant (Ogden et al. 2025). Sockeye spawn in autumn when they lay their eggs in pockets (i.e., redds) in the gravel of streams, and these eggs incubate over the winter, hatching into alevins, before emerging as fry in the spring (Wilson and Peacock 2025). Spawner-to-fry production may be influenced by the number of spawning adults and their condition, the condition of the spawning grounds, and environmental conditions during egg incubation.

During the incubation period, high flows can scour the stream bed, potentially dislodging or burying eggs and alevins, while also increasing fine sediment infiltration, which in turn reduces oxygen supply (DeVries 1997; Ingram 2011). Conversely, if flows are low enough to expose the eggs to air, mortality can occur through desiccation or freezing (Casas-Mulet et al. 2015). While scouring flows during the incubation period can reduce survival, they may have beneficial effects if they occur at other times of year. Over time, fine sediments can accumulate in spawning habitat, filling the spaces between spawning gravels, and causing mortality by reducing dissolved oxygen and trapping emerging alevins (Moring 1982). High-velocity flows, such as those during freshet, can flush fine sediments from spawning gravel, and may be critical to maintaining productive spawning habitat (Reiser et al. 1989).

While desiccation, scour, and sediment flushing occur naturally, they are directly influenced by dam-regulated flow regimes, as is the case in the Okanagan River. Because of this, the Okanagan Basin Implementation Board adopted seasonally varying target flows in 1976, to ensure that flow is suitable for each life stage of salmon in the river (Okanagan Basin Implementation Board 1982), but challenges associated with balancing flood mitigation as well as uncertainty in real-time conditions resulted in flows frequently exceeding or falling below this range (Ng et al. 2025). To address this issue, the Fish/Water Management Tool (FWMT) was developed as a decision support tool that provides real-time predictions of the consequences of water management decisions for fish and other users (Hyatt et al. 2015). Since its implementation in 2004, there have been fewer days in which flows fell below or exceeded the target flow range (Alexander et al. 2024; Ng et al. 2025).

It has been assumed that improved flow compliance has reduced scour and desiccation, thereby improving egg-to-fry survival. However, quantifying this benefit is challenging due to limited fry abundance data prior to FWMT implementation. Ng et al. (2025) found that emerging fry recruitment (i.e., fry per spawner) was higher in the 19 years after the FWMT was implemented, compared to the three years prior. Similarly, Alexander et al. (2024) found a two-fold increase in the ratio of overwintering juveniles (in the fall and winter following emergence) to spawners after brood year 2008 (four years post-FWMT implementation) and concluded that FWMT was a strong contributing factor to the sockeye recovery. Despite this, neither study was able to establish a causal link between fry recruitment and interannual variation in flow rates.

Temperatures experienced during incubation may influence spawner-to-fry production, with embryo survival declining at temperatures of 14-16 ^°^C (Beacham and Murray 1990; Whitney et al. 2013). However, this range is considerably higher than those observed in the Okanagan River, which rarely exceed 10 ^°^C during incubation. Thus, it remains unclear whether interannual variation in incubation temperatures affects survival of Okanagan sockeye.

Beyond the direct effects of incubation conditions, upstream migration conditions may indirectly influence spawner-to-fry production through effects on spawner condition, separate from any effects on abundance due to migration mortality (Fenkes et al. 2016). Elevated river temperatures place considerable stress on migrating sockeye; higher temperatures increase energy use (Rand et al. 2006), promote pathogen development (Crossin et al. 2008), and reduce aerobic scope (Eliason et al. 2011). This results in reduced migration survival when temperatures exceed 18 ^°^C (Martins et al. 2012a), particularly for females (Crossin et al. 2008; Martins et al. 2012b). There is also evidence that high temperatures encountered during migration lead to egg-retention (Quinn et al. 2007), smaller eggs (Braun et al. 2013), reduced time on the spawning grounds (Minke-Martin et al. 2018), and reduced egg-to-fry survival (Patterson et al. 2004), even when individuals successfully reach the spawning grounds and spawn.

Okanagan sockeye begin their 830 km migration up the mainstem of the Columbia River in mid-June, passing nine hydroelectric dams and reaching the confluence of the lower Okanogan River (spelling differs in U.S.A. and we distinguish the Canadian vs. U.S.A. sections in this way) by late June or early July (Fig. S1A). From there, they migrate an additional 185 km up the Okanogan River reaching their spawning grounds above Osoyoos Lake. Summer temperatures in the lower Okanogan River often present a thermal barrier to migration, forcing fish to hold in Lake Pateros at the confluence with the Columbia River and resulting in considerable thermal stress (Miller-Saunders et al. 2024) and en-route mortality (Hyatt et al. 2020). Observations of broodstock collected by the Okanagan Nation Alliance *kł cpəlk stim* hatchery indicate that fish forced to hold in Lake Pateros for an extended period produced fewer eggs compared to fish that were transported past the thermal barrier and held in cooler hatchery waters (pers. comm. D. Stefanovic and T. Marsel, Okanagan Nation Alliance), despite documented negative effects of maternal hatchery holding on egg size (Sopinka et al. 2016). It is therefore plausible that the timing and duration of the thermal barrier may impact both the number and quality of eggs from the fish that survive to reach the spawning grounds.

Here we present a process-based Bayesian model that estimates how flow velocity and temperature, during and preceding the incubation period, influence spawner-to-fry production of sockeye salmon in the Okanagan River. This model allows us to test the following hypotheses: (1) higher pre-spawning flows increase production by flushing fine sediments from spawning gravel; (2) production is reduced in years when incubation-period flows fall outside the FWMT target range (5–28.5 m^3^/s), reflecting increased desiccation and scour risk; (3) interannual variation in incubation temperature has limited effects on production because temperatures during incubation generally remain within ranges suitable for embryo survival; (4) production is reduced in years when the thermal barrier in the lower Okanogan River forms earlier and lasts longer, through carry-over effects of thermal stress on spawner condition. By quantifying the benefits of flow management for salmon and the risks posed by warming temperatures, this work supports the continued application of the Fish/Water Management Tool and provides insight into how similar approaches may support the sustainable management of Okanagan sockeye and other salmon populations.

## Methods

### Study system

#### Salmon enumeration

##### Spawners

Spawner abundance was estimated using a previously published Bayesian model fit to spawner enumeration surveys conducted by the Okanagan Nation Alliance (ONA) in the Okanagan River Index section between Osoyoos and Vaseux lakes (Fig. S1B; Thompson et al. In press). The Index section is the primary spawning grounds for the Osoyoos population, and is the spawning grounds that are sampled by the fry emergence surveys that are used in this study (described below). The Thompson et al. (In press) model accounts for observation error and repeated counts of individuals on the spawning grounds, producing annual posterior distributions of total spawner abundance. For the present analysis, we used the posterior mean and standard deviation of log spawner abundance as inputs to the spawner-to-fry recruitment model (described below), thereby propagating uncertainty from the spawner estimation process.

##### Fry emergence

The ONA has conducted fry emergence surveys in the Okanagan River each spring (March–May) since 2004; trial sampling in 2002 and 2003 was excluded because methods were not standardized. Samples were also excluded in 2018 (brood year 2017) because a different methodology was tried and the peak evening sampling window was not included. Sampling occurs approximately weekly early in the emergence period and increases to every 3–4 days during peak emergence, with reduced frequency thereafter.

Emerging fry are sampled at Vertical Drop Structure 13, which is located on the downstream end of the main sockeye spawning grounds (Index section) for the Osoyoos population (Fig. S1B). Emerging fry are sampled using a fyke net (1 × 2 ft opening; 5 mm stretch mesh) with typically four sets per night, each with a target soak time of 15 minutes. For each set, total fry catch, soak time, and set start time are recorded. Since 2008, a flow meter has been used to estimate the sampled water volume for each set. To reduce heterogeneity in sampling conditions (particularly in early years), our analysis only includes sets conducted between 21:00 and 01:00 with soak times between 5 and 20 minutes, and with sampled volume of at least 10 m^3^.

These surveys were conducted within Syilx title and rights relating to Okanogan sockeye. No field permit was obtained. Animal ethics approval was not obtained for this study. Fish were surveyed using methods intended to minimize stress and mortality, including a blunt cod-end to retain fry alive and shortened soak durations when abundances or flows were high. Fish generally remain alive and in good condition and are released back into the river.

### Environmental covariates

#### Upstream migration thermal barrier

Following Stiff and Thompson (2026), we define the thermal barrier period as the days between June 25 and September 30 when minimum daily water temperatures in the Okanogan River, measured in Malott Washington (USGS station 12447200), exceed 22 ^°^C without a break of at least five days. Daily mean water temperature records were obtained from the U.S. Geological Survey National Water Information System (NWIS) using the dataRetrieval R package (DeCicco et al. 2025). Gaps in the Malott temperature record were filled using a linear mixed-effects model relating daily minimum temperatures at Malott to concurrent temperatures at Oroville (USGS station 12439500) and the Similkameen River (USGS station 12442500), with year-specific random intercepts and slopes for both predictor stations. Although river temperatures can exceed 22 ^°^C before June 25th, we set this as the earliest barrier date because no more than 5% of the run has passed Wells dam (directly downstream) by that date in any of the past 25 years (Columbia River DART, Columbia Basin Research, University of Washington 2025).

The onset date of the thermal barrier is the first day of this period and the duration is the number of days that the thermal barrier lasts. In years in which water temperatures never exceed 22 ^°^C, we define the onset date as 10 days after the latest onset date observed in the dataset. This allowed us to have a continuous covariate for our models, rather than leaving onset date in these years as undefined. Duration is strongly negatively correlated with onset date (*r* = -0.86), reflecting the fact that duration is likely to be longer when onset is early (Fig. 1A-C). To separate these effects, we used residual duration rather than observed duration, calculated from a linear model of duration as a function of onset date, indicating whether a barrier lasted longer or shorter than expected for its onset.

**Figure 1:**
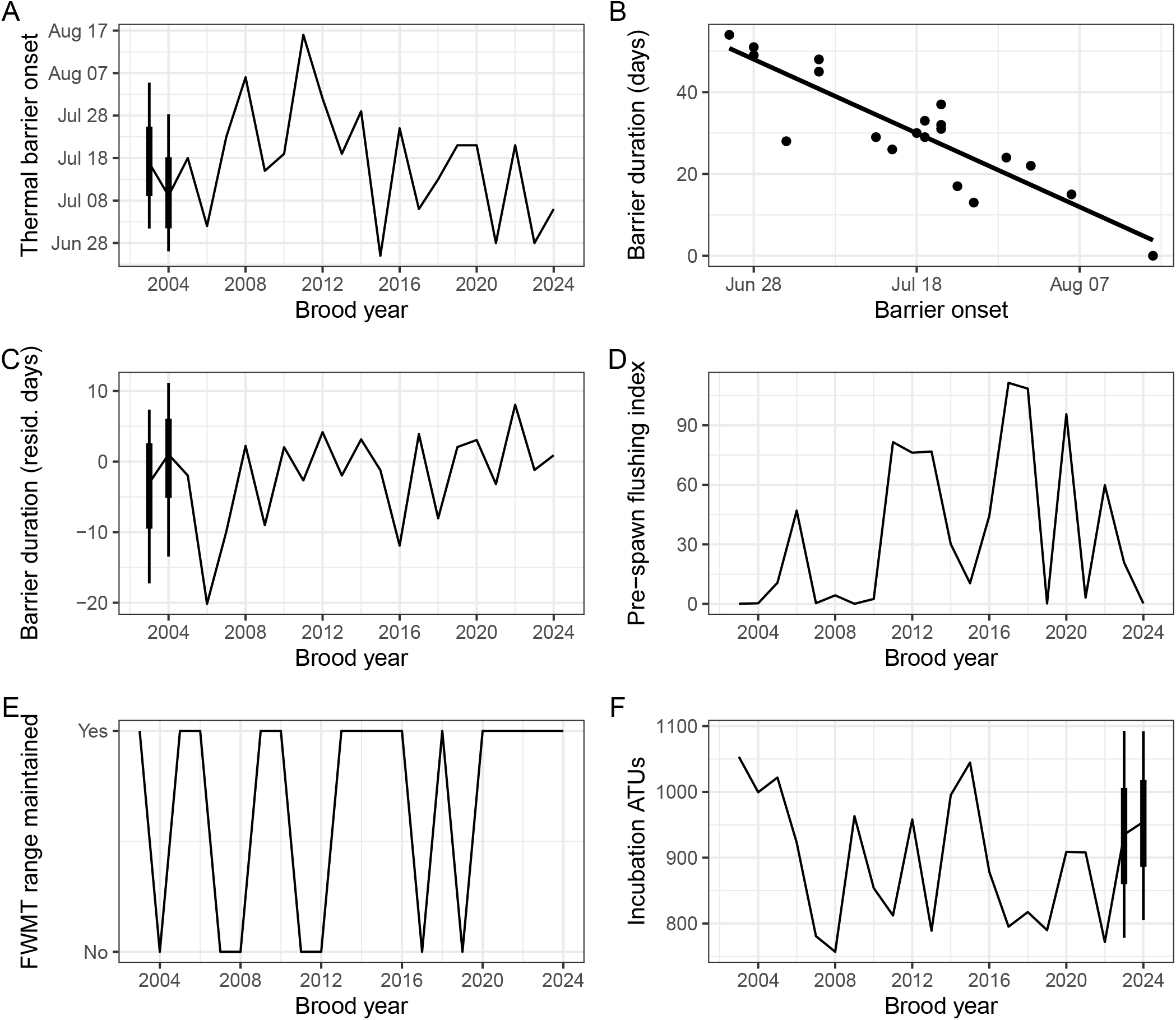
Annual environmental covariates used in the spawner-to-fry production model. (A) Date of the thermal barrier onset in the lower Okanogan River. (B) Linear relationship between thermal barrier onset date and duration (days) used to derive duration residuals. (C) Residual barrier duration (days), after accounting for onset timing. (D) Pre-spawn flushing index. (E) Binary indicator of whether the FWMT range was maintained during incubation. (F) Accumulated thermal units (ATU) during incubation. Environmental data are shown for the brood year affected by each covariate. Lines show observed values or posterior medians in years with imputed data. Intervals (66% and 95%) are shown only for imputed values.

Because water temperatures are not available prior to 2005, we treated thermal-barrier onset and residual duration as partially missing covariates and estimated them within the Bayesian spawner-to-fry recruitment model. For years with missing water temperature data, standardized onset date (mean = 0, SD = 1) was modelled as a linear function of mean June air temperature from the nearby NOAA station USW00094197. The regression coefficients and residual variance were estimated from years with observed onset, and onset dates in missing years were inferred under this fitted relationship and error distribution. Residual duration (scaled by its standard deviation) was also treated as partially missing and was estimated within the Bayesian model by assuming a skew-normal distribution across years; missing values were inferred from this distribution.

#### Pre-spawn flushing flows

We characterized pre-spawn flushing conditions using daily discharge from the Okanagan River near Oliver (Water Survey of Canada hydrometric station 08NM085) during January through September preceding each spawning season (Fig. 1D). To summarize the extent to which flows were capable of mobilizing fine sediments within the spawning gravel, we constructed an annual flushing index:

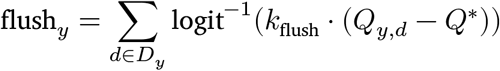

where *Q*_*y,d*_ is the discharge on day *d* in year *y* and *D*_*y*_ includes all days from January through September. The parameters *Q*^*^ and *k*_flush_ define how daily discharge is translated into a flushing intensity between 0 (no mobilization) and 1 (widespread mobilization), representing a gradual transition from negligible gravel movement at low flows to widespread mobilization at high flows. Summing across days allows flushing effects to accumulate over the year preceding spawning.

We selected values for these parameters based on observations of gravel mobility in the Okanagan River spawning grounds, which show that movement begins to occur at around 28 m^3^/s (aligned with the FWMT upper threshold for incubation flows), with full movement occurring at around 70 m^3^/s (Summit Environmental Consultants Ltd. 2002). Thus, we set *Q*^*^ as the discharge corresponding to the approximate midpoint of this transition (50 m^3^/s), and *k*_flush_ to control the steepness of the response so that flushing intensity increases smoothly from near zero to near one across the observed range of gravel movement.

#### Incubation period flow and temperature

We define the incubation period as October 12 to April 10. These dates are defined based on the date that spawners have typically arrived on the spawning grounds (Thompson et al. In press) and just prior to when emerging fry have been most abundant in the emergence samples (Stockwell et al. 2020). To estimate flow rates on the spawning grounds, we use daily flow rates in the Okanogan River near Oliver, which is monitored by the Water Survey of Canada (Hydrometric station 08NM085), and data were extracted directly from the FWMT. This station is directly downstream of the spawning grounds where spawner enumeration and fry emergence sampling occurs (Fig. S1). In each year, we calculate: 1) whether any incubation-day discharge fell outside the FWMT target range of 5–28.5 m^3^/s, and 2) the accumulated thermal units (ATUs) at the end of the incubation period (i.e., April 10; Fig. 1E,F). ATUs were calculated as the cumulative sum of mean daily water temperatures over the incubation period for each brood year.

We used this binary measure of flow impacts because it is a direct assessment of whether FWMT is influencing spawner-to-fry production. We considered using continuous flow metrics that account for duration, frequency, or magnitude, but elected not to because, although they would help with understanding the underlying mechanism, they would not provide an assessment of the FWMT impact, which was a primary goal of this study. Additionally, they would not capture the potential negative effects of both low and high flow events, which the FWMT range is designed to avoid.

### Directed acyclic graph and overall model structure

The assumed causal pathways and observation structure are summarized in a directed acyclic graph (DAG; Fig. 2), which separates (1) annual spawner-to-fry production, (2) within-year emergence phenology linking annual fry abundance to daily emergence intensity, and (3) the observation process linking daily emergence to set-level catches via sampling effort and timing. The DAG guided the structure of our joint Bayesian model, which we describe in the subsequent sections. The model combines multiple submodels in a single joint likelihood implemented in Stan (Stan Development Team 2024) and fit in R version 4.4.3 (R Core Team 2024) using cmdstanr (Gabry et al. 2024).

**Figure 2:**
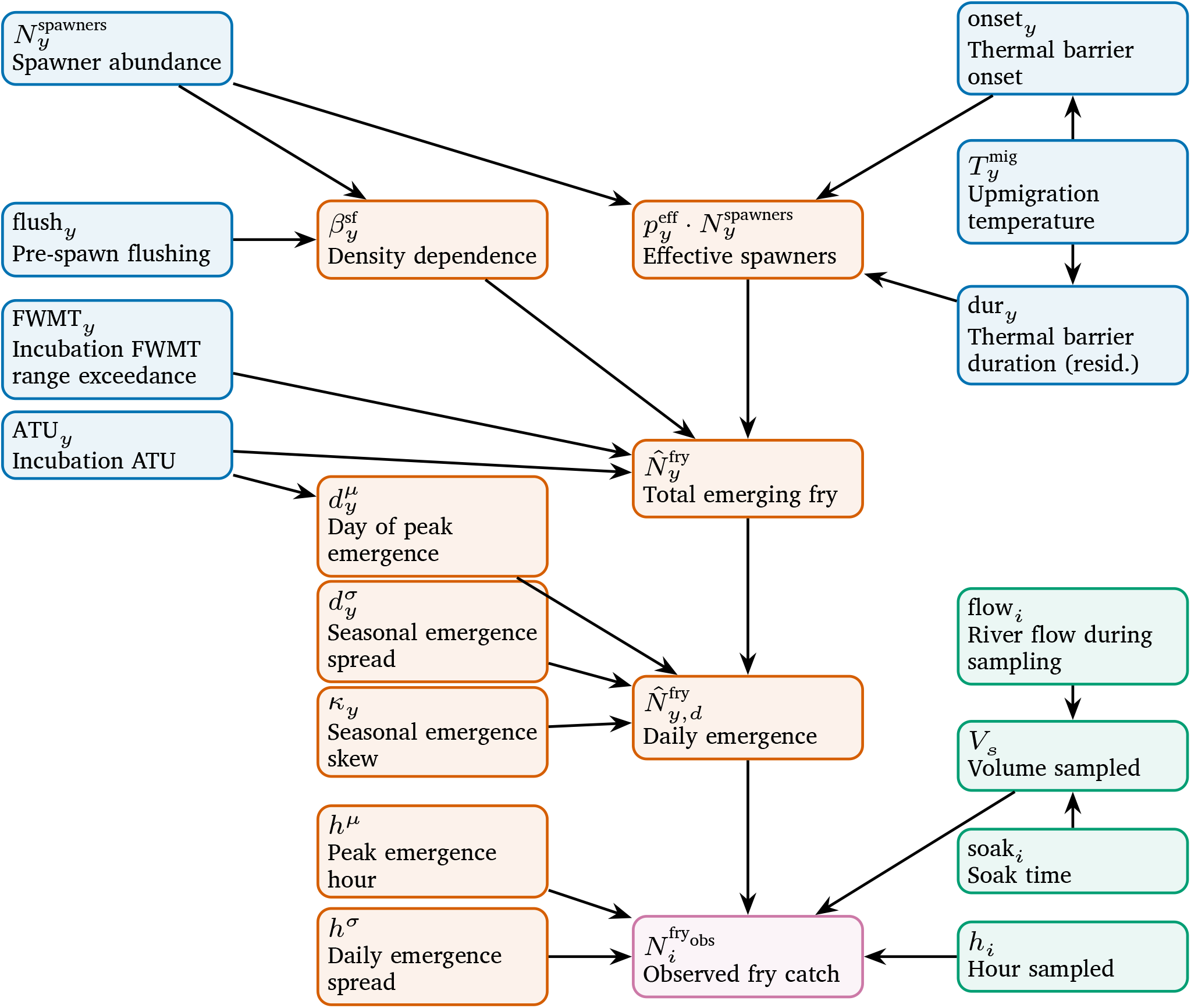
Directed acyclic graph (DAG) illustrating the assumed relationships among spawner abundance, environmental drivers, fry emergence phenology, and observed fry catches at the set level. Arrows indicate conditional dependencies among variables in the hierarchical model. Node colours indicate: environmental and population drivers (blue), latent ecological processes (orange), sampling/observation processes (green), and the observed response (pink).

We used Microsoft 365 Copilot (GPT-5, 2026) as a tool to support Stan model development, including discussion of alternative model structures, generation of example code, and identification of potential causes of convergence issues. All AI-generated suggestions and code were reviewed, tested, and verified by the authors for logic and accuracy before incorporation into the analysis.

### Estimating a metric of yearly fry abundance

The goal of these surveys is to estimate the peak and end dates of emergence. However, they can also provide a relative index of total fry abundance in each year 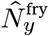. The index is relative because the proportion of all emerging fry that are sampled is unknown. This annual index of fry abundance is treated as a latent state representing the relative seasonal total of emerging fry. It is shared by both the emergence observation submodel and the spawner-to-fry recruitment submodel within a joint likelihood, as outlined in the subsequent sections. The emergence data inform 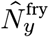, while the recruitment submodel provides its expectation for this index as a function of spawner abundance and environmental covariates.

#### Fry-emergence submodel

To specify the latent emergence process, we modelled fry emergence as a smooth, unimodal skew-normal function of seasonal timing expressed on the day-of-year scale, such that the total index of fry abundance within a given year 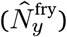 is distributed across the emergence season *D*. We define *D* as the sequence of days between 67 and 154, which corresponds to March 8–June 3 in non-leap years. The total number of fry, as well as the seasonal phenology, vary across years such that:

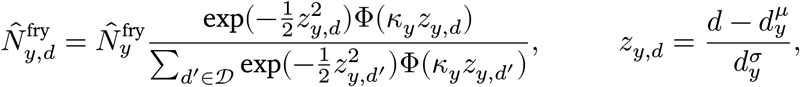

where 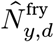 is the expected emergence intensity on day *d* in year *y*. The parameters 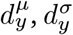, and *k*_*y*_ represent the peak timing, the spread of emergence across days, and the skew of the distribution in year *y*, modelled hierarchically across years (peak and skew on the linear scale; spread on the log scale). We also allowed interannual variation in peak timing to depend on incubation temperature conditions by modelling 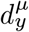 as a linear function of accumulated thermal units (ATUs). The function Φ(.) denotes the standard normal cumulative distribution function. We used standardized (mean = 0, sd = 1) day *d* in the model.

We evaluated alternative formulations of the seasonal emergence phenology, including (i) a model with time-invariant skew *κ*, and (ii) a symmetric Gaussian emergence function. The skew-normal model with interannual variation in skew presented above had substantially improved out-of-sample predictive performance, as quantified using leave-one-out cross validation (ΔELPD > 49).

We linked the latent emergence intensity to the observed fry counts from individual sets using a negative binomial likelihood with an overdispersion parameter *ϕ*^fry^. Thus, the observed fry count in set *i* is modelled as:

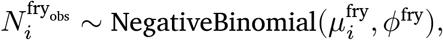

with an expected value

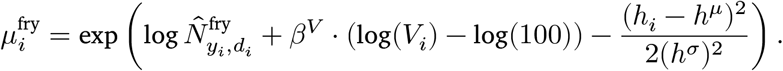

Here, β^*V*^ is the rate at which fry abundance increases with sampled volume *V*_*i*_, expressed relative to 100 m^3^. Within-day variation in emergence is modelled at the observation level using a unimodal Gaussian kernel on standardized log-transformed hours elapsed since 10:00 (*h*_*i*_; sample times range from 21:00 to 01:00). Log-transforming elapsed sampling time captures the observed pattern of fry abundance over the course of the evening (Fig. S3) and substantially improves out-of-sample predictive performance, as quantified using leave-one-out cross validation (ΔELPD = 30.3). The parameter *h*^μ^ is the hour of peak emergence in the day, and *h*^σ^ is the spread of emergence across hours in a day.

Missing sampled volume was imputed jointly within the full Bayesian model using a log-normal regression of sampled volume on soak time and daily flow (both log-transformed). The parameters were estimated based on the observed volumes, and missing values were imputed based on these relationships.

#### Spawner-to-fry recruitment submodel

We modelled expected fry abundance 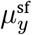 in year *y* as a Shepherd function (Shepherd 1982) of spawner abundance 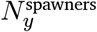:

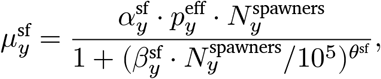

where 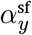 is the density-independent production rate in year *y*, 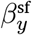 is the rate of density dependence, and θ^sf^ governs the shape of density dependence. 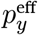 is a proportional scaling factor that reduces the effective reproductive contribution of each spawner, based on thermal conditions experienced during upstream migration (see below). This spawner-to-fry recruitment function reduces to the Beverton-Holt (Beverton and Holt 1957) form when θ^sf^ = 1 and approaches a Ricker (Ricker 1954) form when θ^sf^ → ∞. Thus, it is a generalized version that does not assume a specific shape for density dependence. We divide 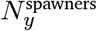 by 10^5^ in the denominator to keep 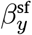 near one, avoiding the need for very small parameter values.

#### Density-independent production during incubation

We allow the density-independent production rate 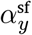 to vary from year-to-year based on environmental conditions experienced during incubation, modelling it as a log-linear function in order to ensure positivity:

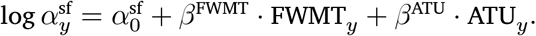

The parameter 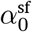is the density-independent production rate during average conditions and the β coefficients are the slope parameters that estimate the effect of accumulated thermal units during incubation (β^ATU^) and of whether or not discharge exceeded the FWMT target range at any point during the incubation period (β^FWMT^).

#### Thermal barrier impacts on per-capita spawner effectiveness

We model the effect of thermal conditions during upstream migration on spawner-to-fry production as a reduction in per-capita spawner reproductive effectiveness. We use a logistic function to represent this effect, reflecting the assumption that delayed thermal-barrier onset can increase effectiveness only up to the point at which all returning salmon have successfully migrated, beyond which additional delays confer no further benefit:

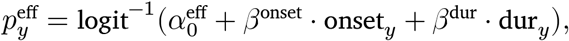

where 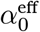 represents baseline per-capita spawner effectiveness on the logit scale under average conditions, β^onset^ describes the effect of thermal-barrier onset date, and β^dur^ is the additional effect of barrier duration beyond that explained by onset timing.

#### Density dependence and pre-spawn flushing

We assume that density dependence during spawner-to-fry recruitment occurs primarily through crowding on the spawning grounds. As spawner abundance increases, there is greater competition for suitable spawning sites. This pushes spawners onto suboptimal gravel and can lead to superimposition, whereby later-arriving spawners build their redds on top of those built by earlier-arriving fish, reducing egg survival through disturbance and displacement (Burgner et al. 1969; Essington et al. 2000). Thus, we assume that the effects of pre-spawn flushing of fine sediments should impact the population by changing the amount and quality of available spawning gravel (Moring 1982; Reiser et al. 1989).

We modelled annual variation in density dependence as a function of the flushing index defined above. Specifically, we allowed the density-dependence parameter to vary log-linearly with the standardized flushing index:

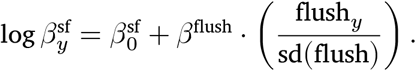

#### Process error and spawner uncertainty

We assume that the annual fry abundance index 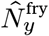 deviates from its expected value 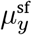 due to process error 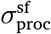, with log abundance following a normal distribution around the expected value:

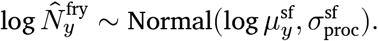

To propagate uncertainty in spawner abundance from our prior model (i.e., Thompson et al. In press), we model log spawner abundance log 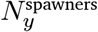 as a latent variable:

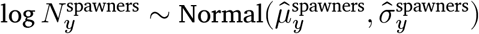

where 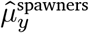 and 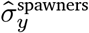 are the mean and standard deviation of log spawner abundance in a given year in that model. Full prior specifications for all parameters in the model are provided in Table S1.

### Quantifying environmental impacts on spawner-to-fry production

We estimated scenario-based contrasts in predicted fry production by varying each environmental covariate while holding all others at their mean value. Except when estimating its effect, the FWMT incubation flow rate range was assumed to be maintained. Predictions were generated at the median observed spawner abundance over the study period. For each contrast, fry production was estimated with the focal variable set to the 15th (low or short) and 85th (high or long) percentiles of its observed distribution. For the FWMT effect, we compared scenarios where incubation flows were maintained within the FWMT range (i.e., FWMT_*y*_ = 0) throughout the incubation period and where the flow exceeded this range at least once during incubation (i.e., FWMT_*y*_ = 1). For each posterior draw, we calculated the percent difference between scenarios. Percent change was calculated using the scenario with higher median predicted fry production as the reference state, such that all estimates are expressed as reductions in spawner-to-fry production.

### MCMC Chains

We ran four parallel Markov chain Monte Carlo (MCMC) chains for 2000 iterations each, discarding the first 1000 as warm-up. We assessed convergence using the rank-normalized version of the Gelman-Rubin diagnostic (Vehtari et al. 2021), and all parameters showed good convergence 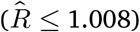, with 100% of 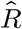 values below 1.01.

## Results

### Fry emergence

Fry emergence abundance and timing varied among brood years (Fig. 3). The fry abundance index ranged from 558 in brood year 2017 to 15 705 in brood year 2009 (Fig. 3B). Peak emergence timing 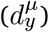 ranged from April 7 in brood year 2015 to May 4 in brood year 2010 (Fig. 3C). Peak emergence timing 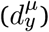 occurred earlier in years with higher accumulated thermal units during the incubation period (Fig. S4), with full posterior support for a negative relationship.

**Figure 3:**
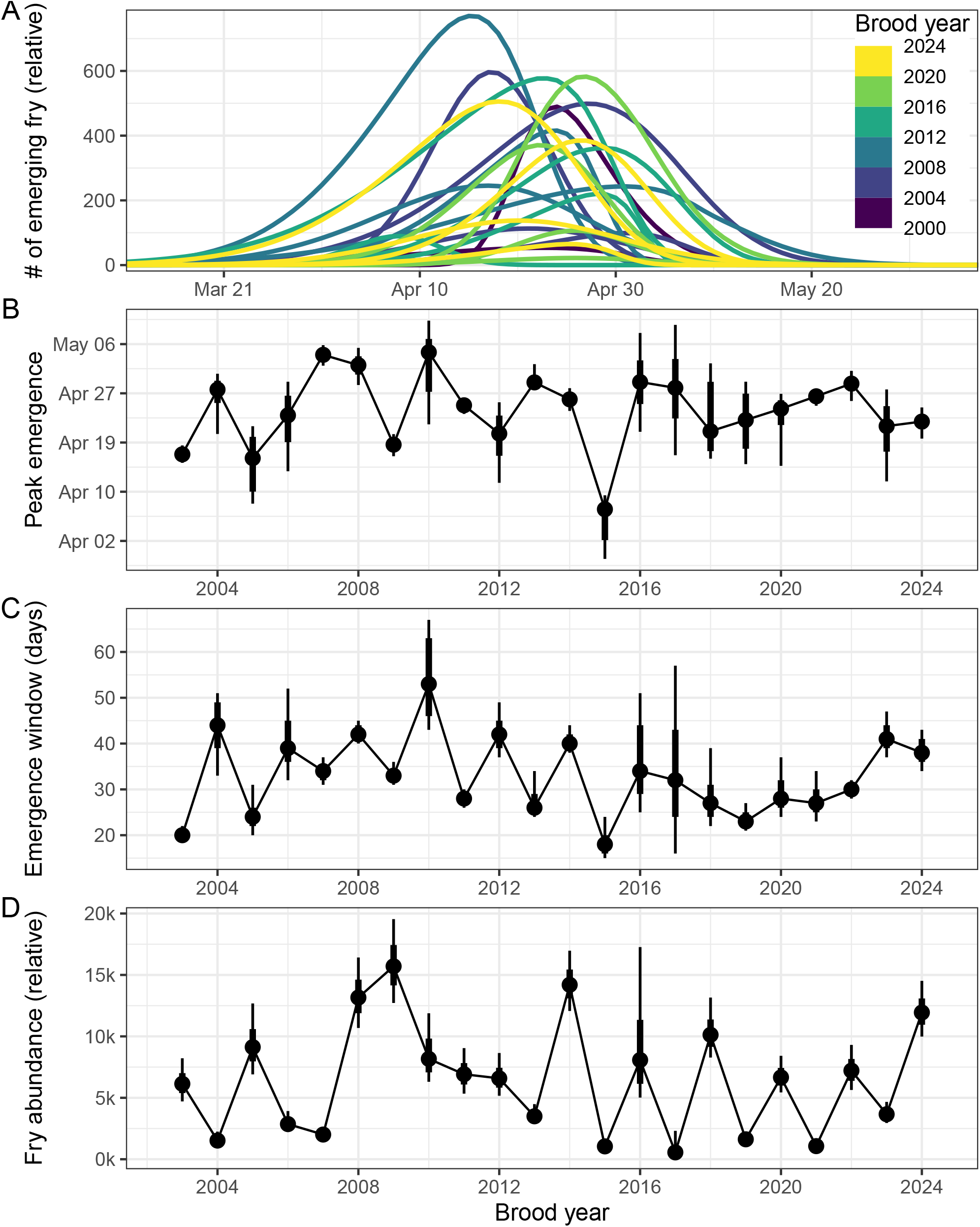
Estimated fry emergence timing and relative abundance from the fry-emergence submodel. (A) The estimated relative abundance at emerging fry across dates, where each line represents a single year coloured by 4-year block. (B) Estimated peak fry-emergence day by year. (C) Estimated width of the emergence window in days by year. (D) Estimated relative fry abundance index by year. Points and error bars in panels B through D show the posterior medians with 66% and 95% credible intervals. Note that brood year 2017 emergence samples were not included; estimates for that year were obtained through model-based imputation informed by the fitted hierarchical model and other years of data.

The width of the emergence window (i.e., number of days over which 95% of emergence occurred) varied from as short as 18 days in brood year 2015 to as long as 53 days in brood year 2010 (Fig. 3D). The parameters describing emergence timing were correlated. In years with later peak timing 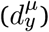, emergence windows tended to be broader (r = 0.64, 95% CrI: 0.22 to 0.85) while skew tended to decrease with later peak timing (r = -0.45, 95% CrI: -0.76 to 0.07; Fig. 3A).

### Drivers of spawner-to-fry production

Relative fry abundance increased with spawner abundance in a density-dependent manner (Fig. 4A). The shape of this recruitment relationship was consistent with Beverton-Holt type compensation, with no evidence of overcompensation (i.e., θ^sf^ ∼ 1; Fig. S5). There were substantial yearly deviations from the average recruitment relationship (Fig. 4A), with thermal barrier dynamics and river discharge during incubation identified as the strongest drivers (Fig. 4B). This is illustrated in Fig. 4A by the fitted lines, which show the estimated recruitment relationships for early (yellow; June 29) and late (purple; July 29) thermal barrier onsets and for incubation flows that remained within (solid) or fell outside (dotted) the FWMT target range.

**Figure 4:**
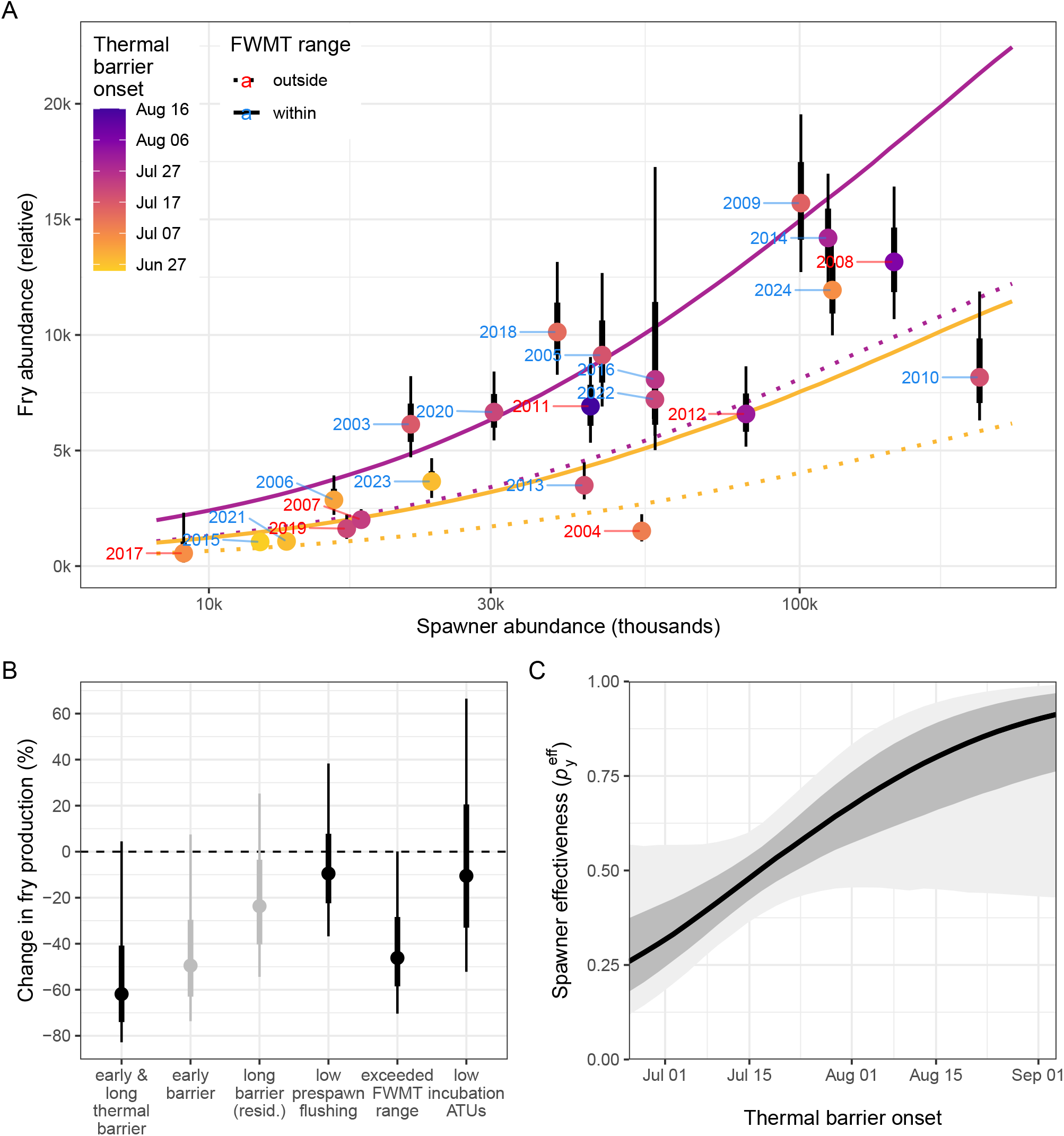
Estimates from the spawner-to-fry recruitment submodel. (A) Estimated relationship between spawner abundance (log axis) and emerging fry abundance (relative index). Points and their error bars show annual posterior estimates of fry abundance coloured by thermal barrier onset date; brood-year labels are coloured by whether or not incubation flows were within the FWMT range. The lines show posterior median spawner–fry curves under two thermal barrier onset scenarios (early: Jun 29; late: Jul 29) and two FWMT scenarios (solid = within FWMT range (5–28.5 m^3^/s), dotted = outside). (B) Estimated percent change in fry production associated with contrasts in environmental conditions and flow management. For continuous variables, contrasts are based on the 15th and 85th percentiles of the observed data. Black points indicate primary contrasts, while gray points indicate the component effects used to derive the combined thermal barrier contrast. (C) Posterior relationship between thermal barrier onset date and per-capita spawner effectiveness. Points and lines are posterior medians; shaded areas and error bars show 66% and 95% credible intervals.

When the thermal barrier begins early (June 29) and persists for a long duration, fry production was estimated to change by approximately -62% (95% CrI: -83% to 4%) relative to years with late onset (July 29) and short duration, with a 97% probability of a negative effect (Fig. 4C). This combined effect was larger than the estimated effects of any other environmental driver and was primarily the result of variation in barrier onset date. When the thermal barrier begins early, fry production was estimated to change by approximately -49% relative to a late onset (95% CrI: -74% to 7%), with a 96% probability of a negative effect. After accounting for variation in onset date, the estimated effect of duration was weaker, with long versus short thermal barrier duration changing fry production by -24% (95% CrI: -54% to 25%), with a 86% probability of a negative effect.

Fry production in years with low pre-spawn flushing was estimated to differ by -9% relative to years with high flushing (95% CrI: -37% to 38%), with a 73% probability of a negative effect. When flows during the incubation period fall outside of the target range from the FWMT (5–28.5 m^3^/s), fry production was estimated to change by -46% (95% CrI: -70% to 0%), with a 97% probability of a negative effect.

In years with low incubation accumulated thermal units, fry production was estimated to differ by -11% relative to years with high incubation accumulated thermal units (95% CrI: -52% to 66%), with a 64% probability of a negative effect.

## Discussion

Our analysis shows that spawner-to-fry production is strongly influenced by thermal barrier dynamics and whether flows during incubation remain within the FWMT target range. In contrast, neither pre-spawning flows nor incubation temperature meaningfully influence production, despite the strong influence of temperature on emergence timing. These results are consistent with thermal barrier dynamics influencing production through carry-over effects on spawner condition, and with incubation flow conditions influencing egg and embryo survival through scouring and desiccation. These findings highlight the growing risk posed by climate warming during the upstream migration stage through its effects on thermal barrier dynamics. They also provide evidence that managing flows using the FWMT benefits fry production, suggesting that effective flow management may partially offset climate-related impacts on freshwater production.

While the effects of the Okanogan River thermal barrier on migration delay and survival of returning sockeye are well documented (Hyatt et al. 2020; Murauskas et al. 2021), our analysis suggests that it is also a major driver of spawner-to-fry production. We estimate that production is approximately 62% lower in years with early-onset, long-duration thermal barriers than in years with late-onset, short-duration barriers. This combined effect was driven primarily by variation in barrier onset date, although longer barrier duration contributed additional reductions in production after accounting for onset. These impacts likely arise from the physiological stress associated with holding and migrating through warm water (Eliason et al. 2011; Fenkes et al. 2016), representing a carry-over effect from the adult migration stage to fry production (Gosselin et al. 2021). This is consistent with the observations from the Okanagan Nation Alliance hatchery, where reduced egg mass and egg number were found in fish that experienced prolonged holding below the thermal barrier (pers. comm. D. Stefanovic and T. Marsel, Okanagan Nation Alliance). Our analysis does not allow us to distinguish which aspects of spawner condition are affected, but previous work suggests that elevated temperatures can influence reproductive success through increased egg retention (Quinn et al. 2007), increased pathogen development (Crossin et al. 2008), and reduced egg-to-fry survival (Patterson et al. 2004).

Although the thermal barrier in the Okanogan River has been documented since the 1960s (Murauskas et al. 2021), it now occurs more frequently, with earlier onset and longer duration (Hyatt et al. 2020), and is projected to intensify under climate change, shifting from an episodic occurrence to a chronic constraint on migration (Stiff and Thompson 2026). As a result, the Okanogan River thermal barrier represents a key bottleneck in the life cycle that may pose a risk to the long-term viability of Okanagan sockeye.

Our analysis finds that spawner-to-fry production is higher when Okanagan River flows remain within the FWMT target range of 5–28.5 m^3^/s during the incubation period. Production was estimated to be approximately 46% lower in years when flows fell outside the target range, corresponding to an increase of approximately 86% when flows remained within it. This result provides quantitative support for the effectiveness of the FWMT, consistent with its design to minimize mortality from both scour at high discharge and desiccation at low discharge (Hyatt et al. 2015). Previous studies documented increases in fry and pre-smolt production following FWMT implementation (Alexander et al. 2024; Ng et al. 2025), but were unable to link these changes directly to interannual variation in flow conditions. Our results suggest that improved compliance with the target flow range is the likely mechanism underlying these patterns, underscoring the value of flow management as a tool for supporting freshwater production.

In contrast, we find limited support for the hypotheses that pre-spawning flows and incubation temperature influence spawner-to-fry production. The limited support for effects of pre-spawning flows may indicate that there is a relatively small influence of sediment flushing on fry production in this system. Alternatively, the flushing index used here may not adequately capture the relationship between discharge and sediment mobilization. However, because it is based on empirical observations of gravel mobilization in the Okanagan River (Summit Environmental Consultants Ltd. 2002), this seems unlikely. The limited support for effects of incubation temperature is consistent with previous evidence showing that negative impacts of temperature on embryo survival only begin to occur when temperatures are higher than those observed in the Okanagan River (Beacham and Murray 1990; Whitney et al. 2013).

The structure of our model was guided by an explicit causal framework represented in our DAG (Fig. 2), which formalizes the assumed links between observed data and underlying processes, and provides a basis for evaluating causal relationships (Pearl et al. 2016; McElreath 2020). Nevertheless, the validity of our inferences depends on the accuracy of these assumptions. In particular, we assume that our index of relative fry abundance provides a reasonable proxy for absolute fry abundance. We also assume that fry emergence can be adequately described using a skew-normal distribution over the season and a lognormal distribution within each evening. More generally, all environmental covariates in the model are represented using indices that approximate underlying processes rather than directly measuring them. For example, thermal barrier dynamics are represented using a 22 ^°^C threshold over the period from June 25 to September 30, which is unlikely to fully capture the thermal exposure experienced by migrating fish. Finally, the relationships specified in the DAG are implemented using specific functional forms, including Shepherd recruitment, which may not fully capture the true underlying processes.

These results provide important insight into the potential causes of the Okanagan sockeye salmon recovery by offering quantitative support for previously hypothesized links between environmental conditions, management actions, and spawner-to-fry production (Alexander et al. 2024). In particular, our analysis supports the hypothesis that FWMT compliance has been an important contributor to the recovery. However, our results demonstrate the benefits of the FWMT only at a single life stage. To assess how these benefits persist through the sockeye life cycle and influence adult recruitment, integration into full life-cycle models will be necessary. Despite this, our results suggest that management measures such as FWMT may help to offset some factors that negatively impact survival, including dam passage-related mortality, the Okanogan River thermal barrier, and marine heat waves. Many of these stressors are expected to be exacerbated by climate change and are not easily managed (Thorstad et al. 2021; Connors et al. 2025). In this context, by providing a quantitative assessment of FWMT effectiveness, our model also establishes a framework for reassessing its performance as additional data become available.

Although this analysis is grounded in the Okanagan system and the specific context of the FWMT, the underlying mechanisms have broader relevance for other salmon populations. Our findings highlight the benefits of limiting scour and desiccation during the incubation life stage; in systems where flows are actively managed, adopting fish-friendly flow regimes would likely provide similar improvements in production. Even in systems without direct flow regulation, habitat restoration measures that reduce the frequency or intensity of extreme flow conditions may also increase fry production. Additionally, return migration temperatures are already a concern in other watersheds, even where temperatures do not consistently reach levels that fully impede migration (e.g., Fraser and Somass Rivers; Eliason et al. 2011; Brown et al. 2026). The impacts of the thermal barrier observed in the Okanagan River may therefore provide insight into the consequences of continued warming in these and other systems. Overall, these findings illustrate how targeted management measures such as the FWMT can help mitigate the negative effects of extreme environmental conditions, while potentially buffering salmon populations against some of the impacts of climate change.

## Acknowledgements

This work stands on the shoulders of Okanagan Nation elders, leaders, knowledge keepers, youth, and others who have desired for over 100 years to re-establish a balanced ecosystem approach to water management for salmon. To the present-day, Syilx Okanagan community members have prioritized being on the land and water to measure and observe the results of FWMT. This story is part of an endeavour to better understand our relationship to place and value in community, culture, history, and the larger vision of the Okanagan Nation Elders to heal the river by *kt cp’əlk’ stim’* (“bringing it back”).

We thank the ONA field crews for their dedication in collecting the spawner and fry data that made this study possible. We also thank the Canadian Okanagan Basin Technical Working Group (COBTWG) which includes participants from the ONA, DFO, and the Province of British Columbia Ministry of Water, Land, and Resource Stewardship. We are also grateful to the late Kim Hyatt who was instrumental in establishing the field surveys and in developing the Fish/Water Management Tool.

We used Microsoft Copilot (GPT-5, 2026) to proofread portions of the manuscript (e.g., mathematical notation and description of the model), focusing on grammar, clarity, and accuracy. All AI-generated suggestions were assessed by the authors and verified for accuracy before being incorporated. The authors take full responsibility for the analyses, interpretations, and conclusions presented in the manuscript.

## Author contributions

Patrick L. Thompson: Conceptualization, Methodology, Formal Analysis, Software, Validation, Visualization, Writing – Original Draft, Review & Editing, Project Administration.

Karilyn Alex: Conceptualization, Investigation, Data curation, Funding acquisition, Writing - Review & Editing; Howard Stiff: Conceptualization, Data curation, Writing - Review & Editing; Athena Ogden: Conceptualization, Data curation, Writing - Review & Editing;

## Funding

The research was supported by a collaborative agreement between Fisheries and Oceans Canada (DFO) and the Okanagan Nation Alliance (ONA) to assess the cumulative impacts of water management, habitat disturbances, and climate change on production variation of Osoyoos Lake Sockeye salmon. The ONA receives funding from the Douglas County Public Utility District.

## Competing Interests

Douglas County Public Utility District provides funding to the ONA for the ongoing deployment of the Fish Water Management Tool as well as monitoring and data synthesis regarding its impacts on fish in the Okanagan basin. The Fish Water Management Tool is among the factors examined in this study. The funder had no role in the analysis, interpretation of results, preparation of the manuscript, or decision to publish.

## Data Availability

The data used in this study are the property of the Okanagan Nation Alliance and are subject to a data sharing agreement. Please contact them directly concerning availability and acknowledgements: Sylix.org.

All scripts and stan model code are available for public access on GitHub (https://github.com/SiRE-P/okanagan_spawn_fry_model) and Zenodo (10.5281/zenodo.22968425).

## Supplementary Materials

### Priors

**Table S1:**
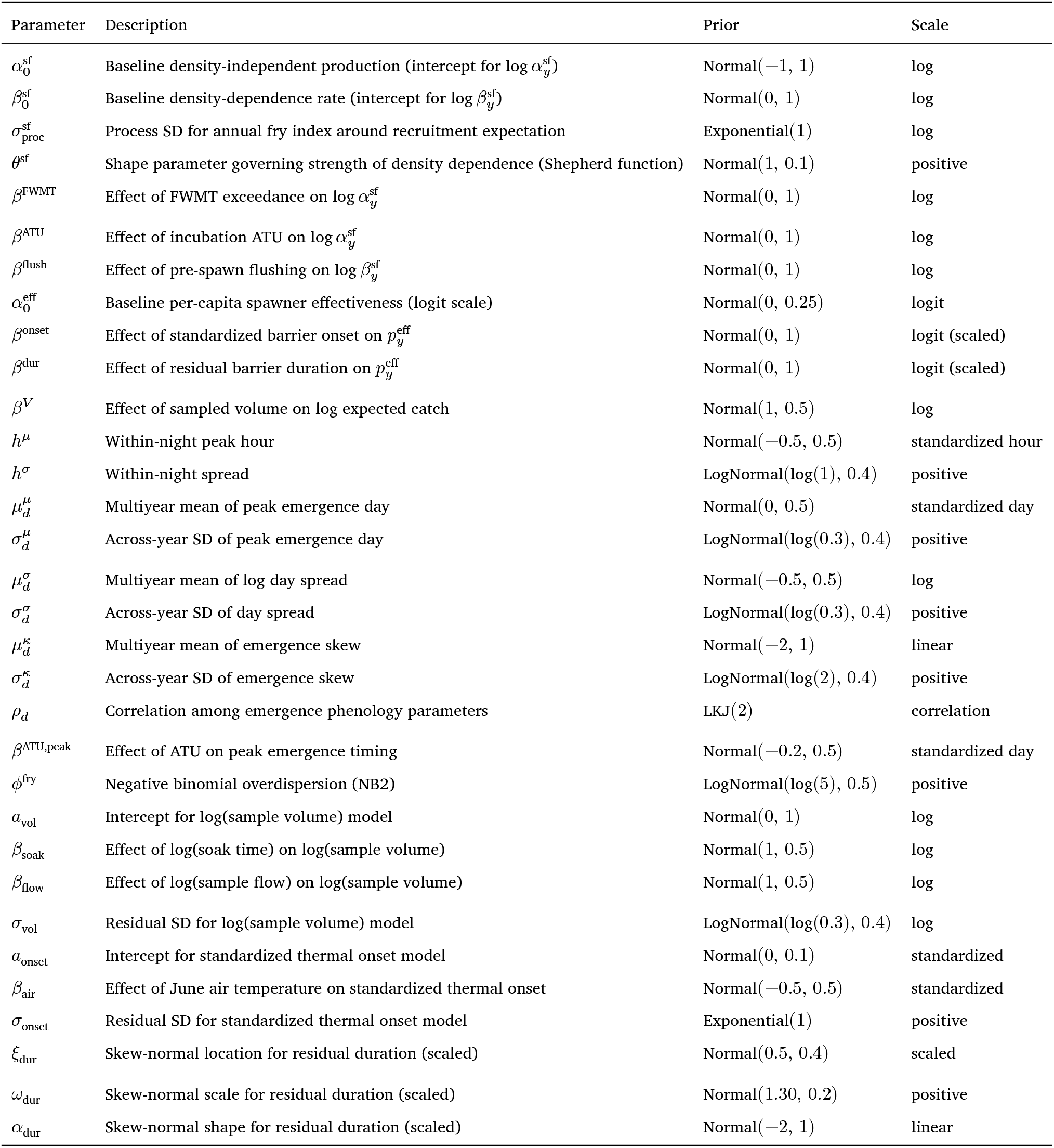
Prior distributions for all parameters in the joint Bayesian model.

### Supplementary Figures

**Figure S1:**
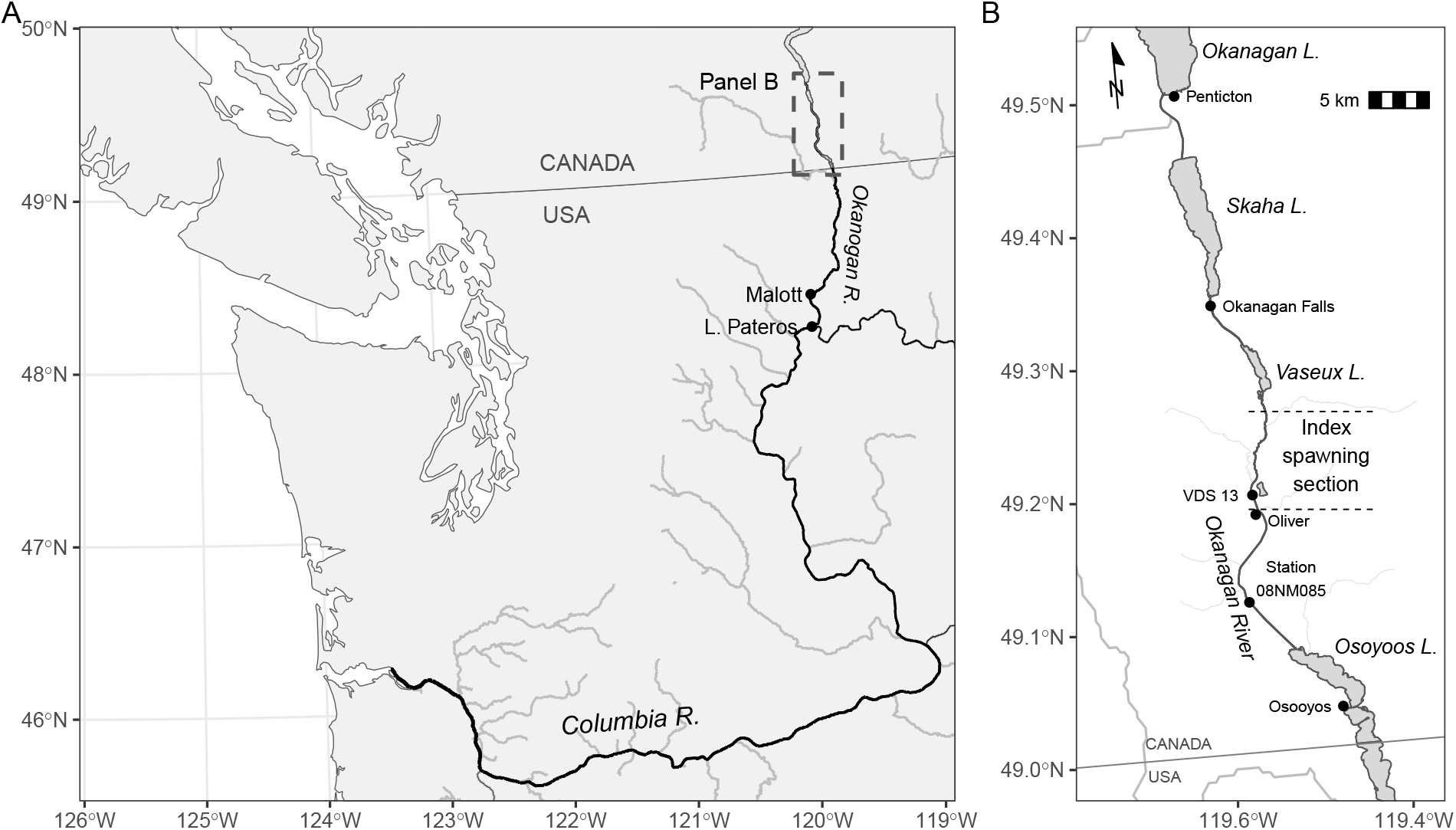
Map of the Columbia and Okanagan/Okanogan River basins. Panel A shows the broader Columbia River Basin downstream of the Okanagan with the mainstem and Okanogan Rivers in black and all tributaries of stream order five or lower in grey. Panel B shows a portion of the Okanagan Basin that covers the index (main) spawning grounds for Osoyoos sockeye in the Okanagan River. The mainstem Okanagan River is in black and other rivers and tributaries of order five and size are shown in grey. Station 08NM085 is the Water Survey of Canada hydrometric monitoring station. VDS 13 is the vertical drop structure where the fry emergence survey is conducted. Spatial data are from GeoBC (Province of BC), HydroRIVERS (Lehner and Grill 2013), and the rNaturalEarth package (Massicotte and South 2026). Map projection: NAD83 / BC Albers (EPSG:3005).

**Figure S2:**
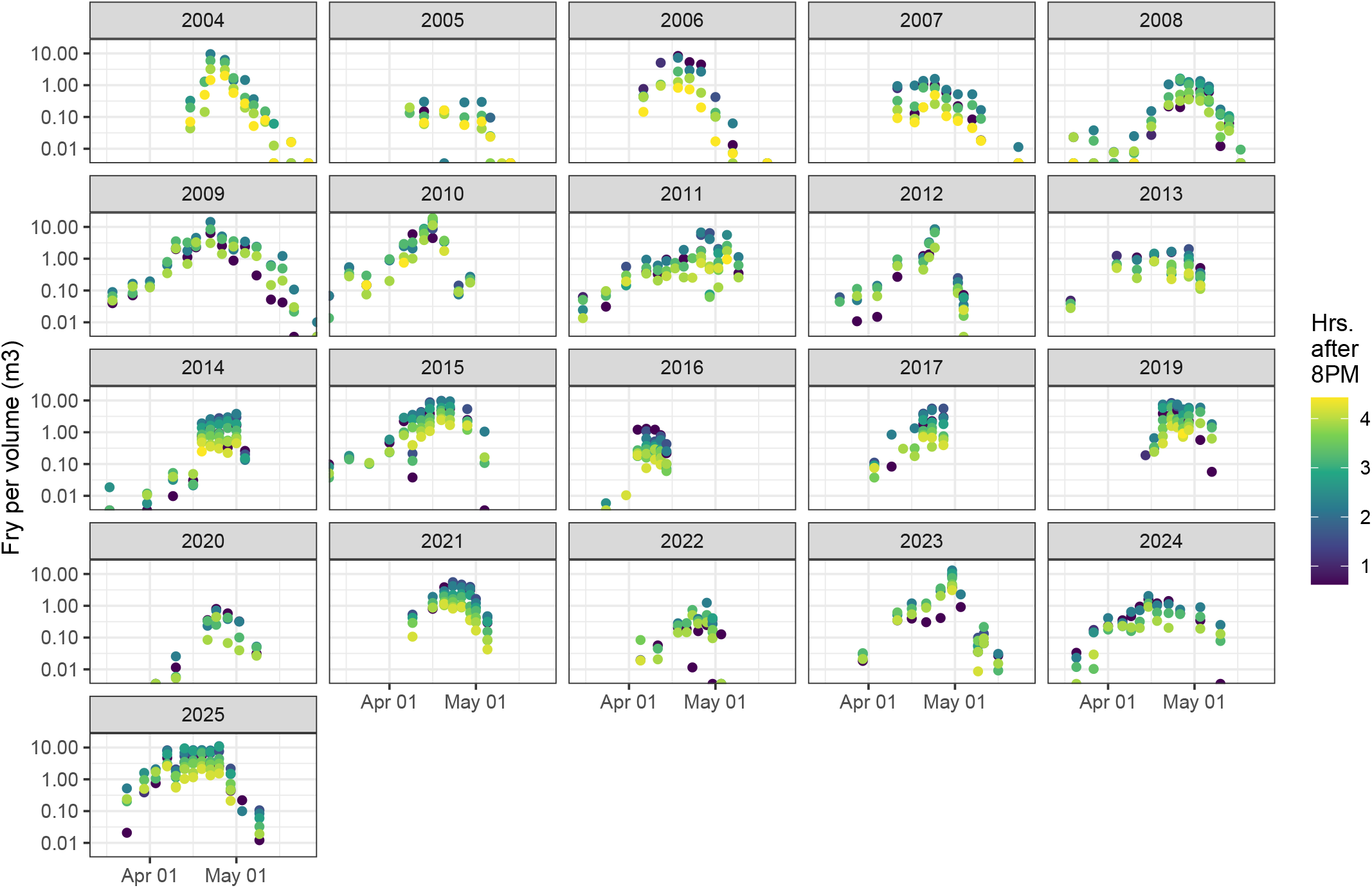
Observed fry per volume in sets over days in each year (i.e., brood year + 1). Colours show the hour that each set was taken.

**Figure S3:**
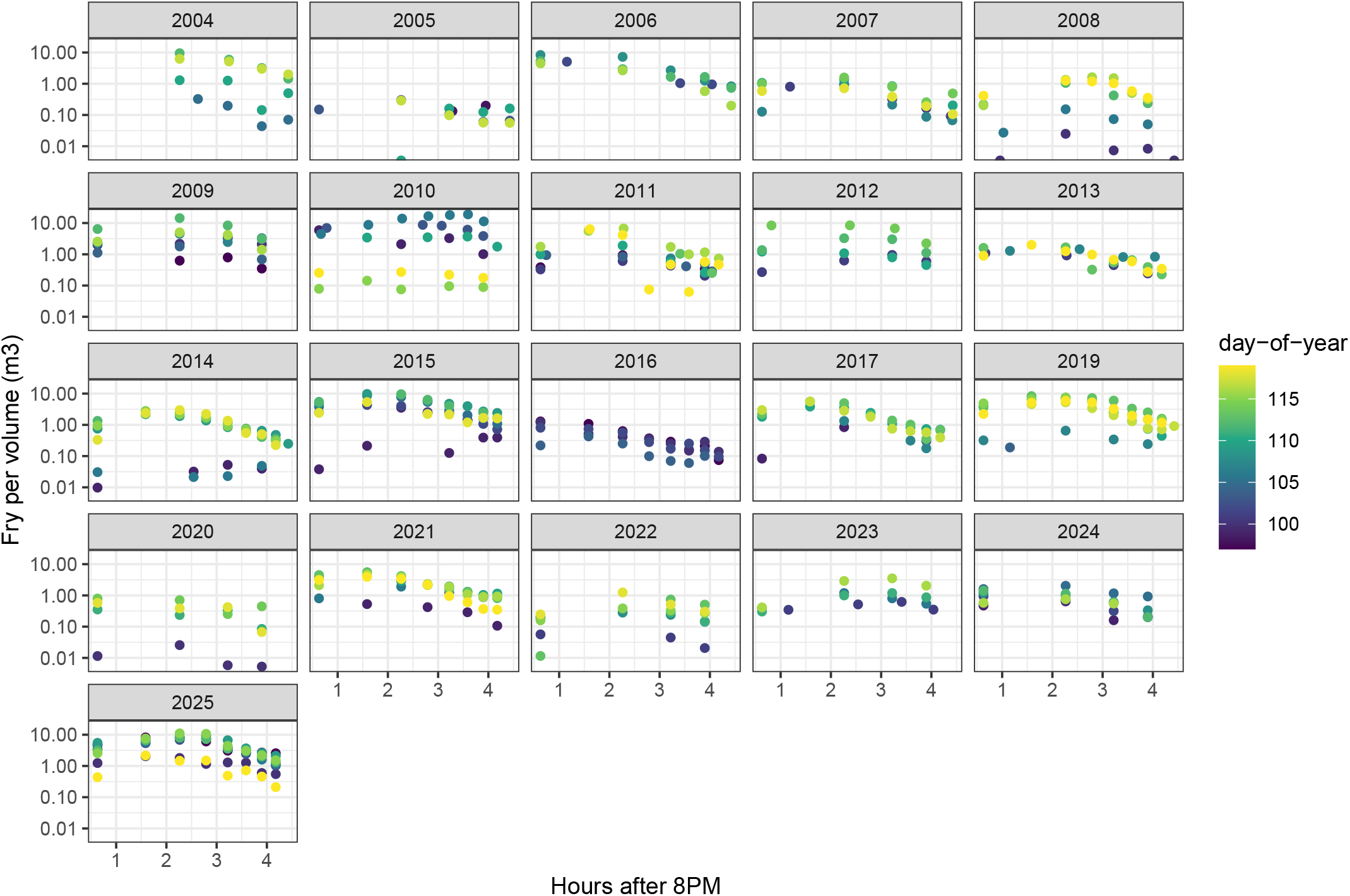
Observed fry per volume of water sampled over hours elapsed since 20:00. Sets are restricted to those between day-of-year 96 and 120, for visual clarity. Years are sample year (i.e., brood year + 1).

**Figure S4:**
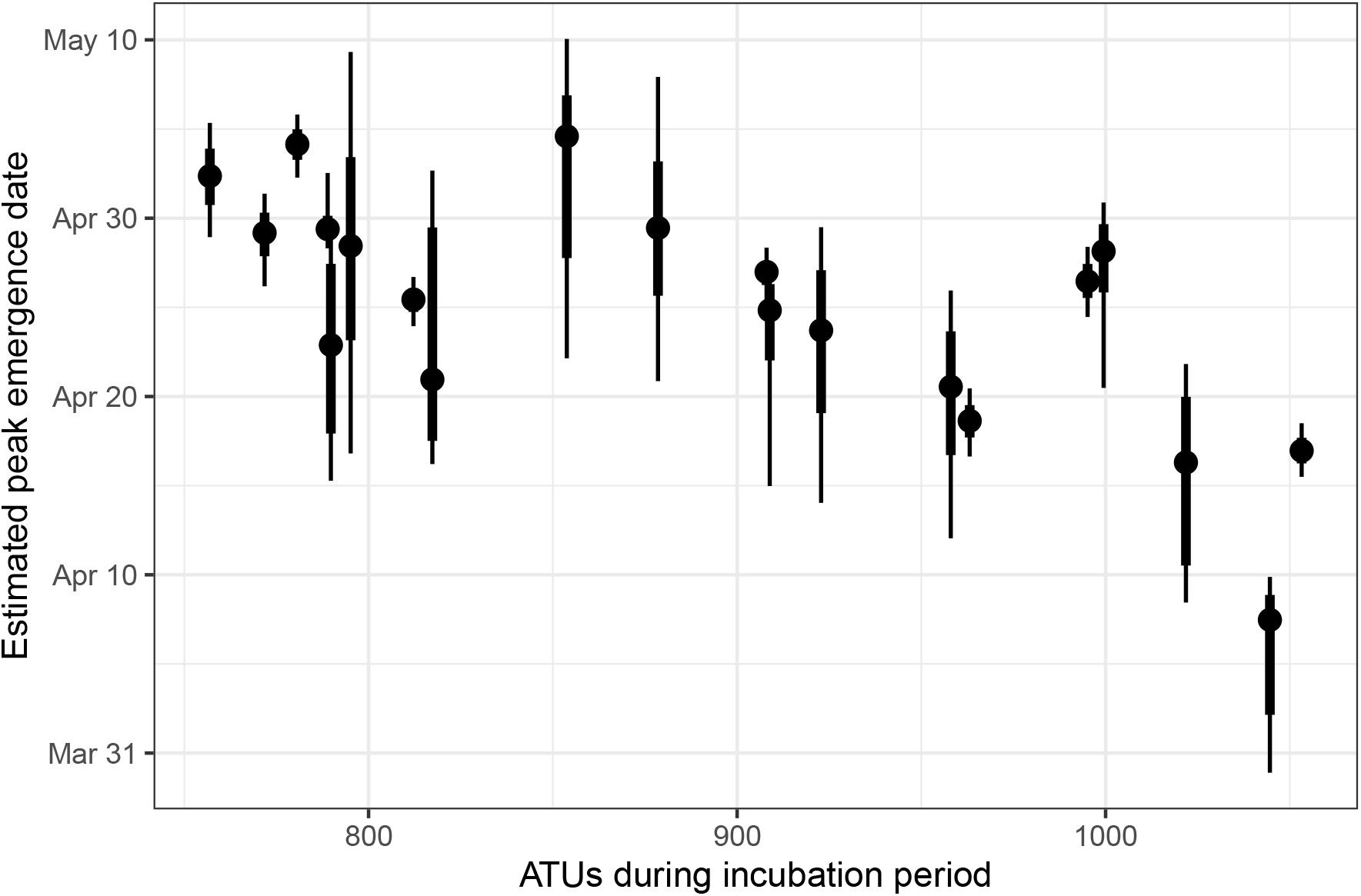
Date of peak emergence timing vs. accumulated thermal units (ATUs) during the incubation period.

**Figure S5:**
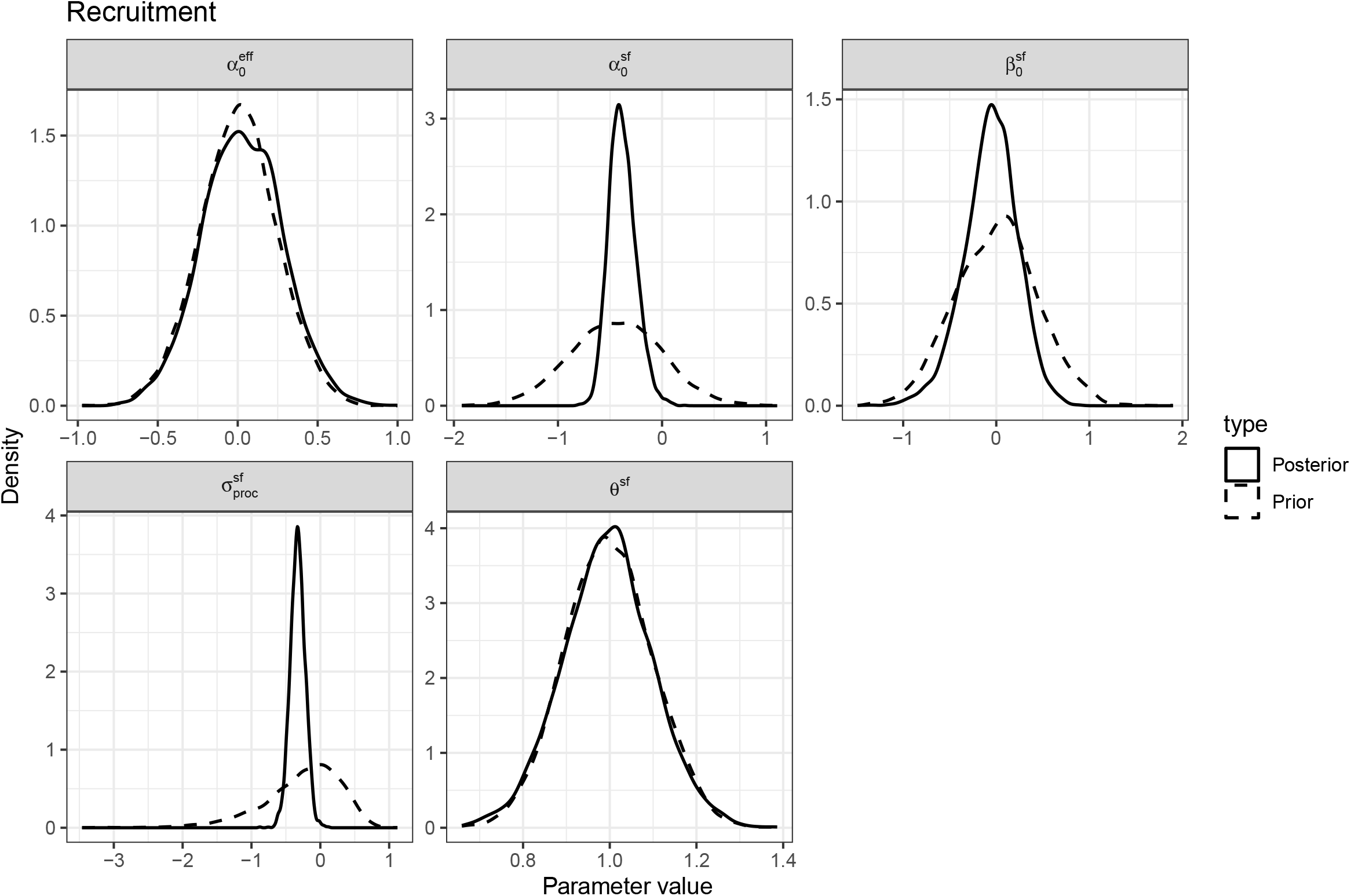
Comparison of prior and posterior for recruitment parameters.

**Figure S6:**
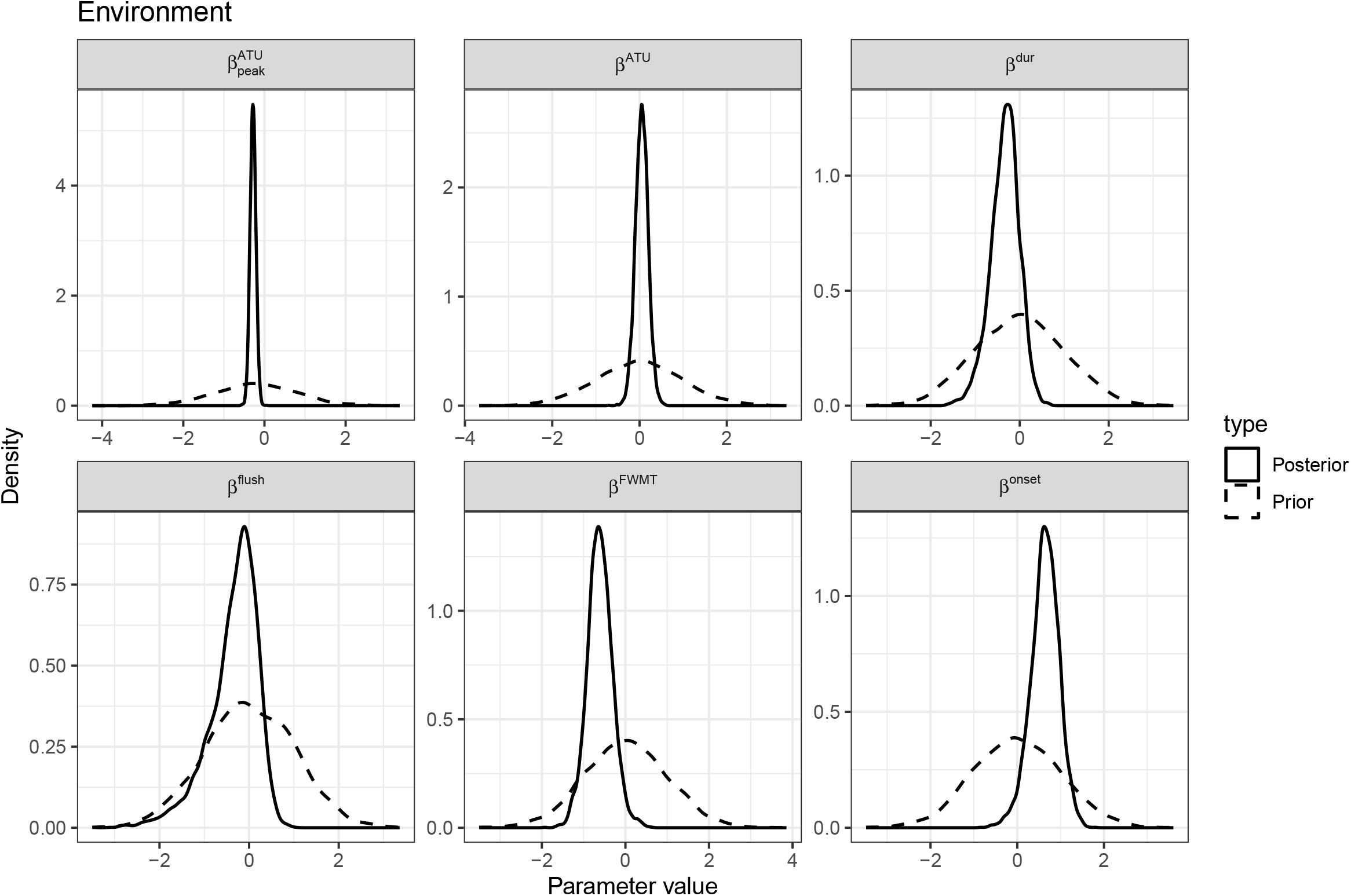
Comparison of prior and posterior for environmental effect parameters.

**Figure S7:**
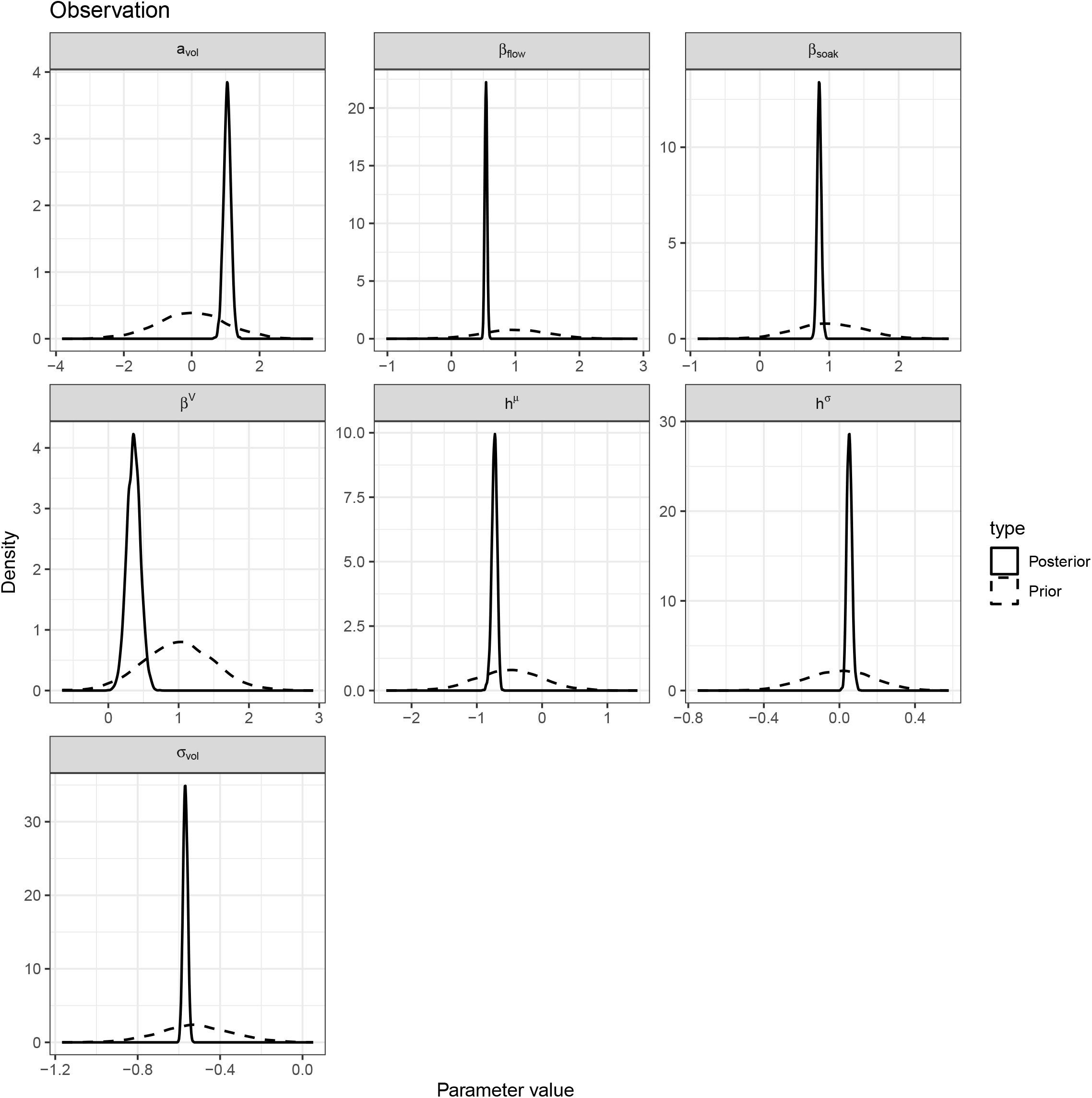
Comparison of prior and posterior for observation parameters.

**Figure S8:**
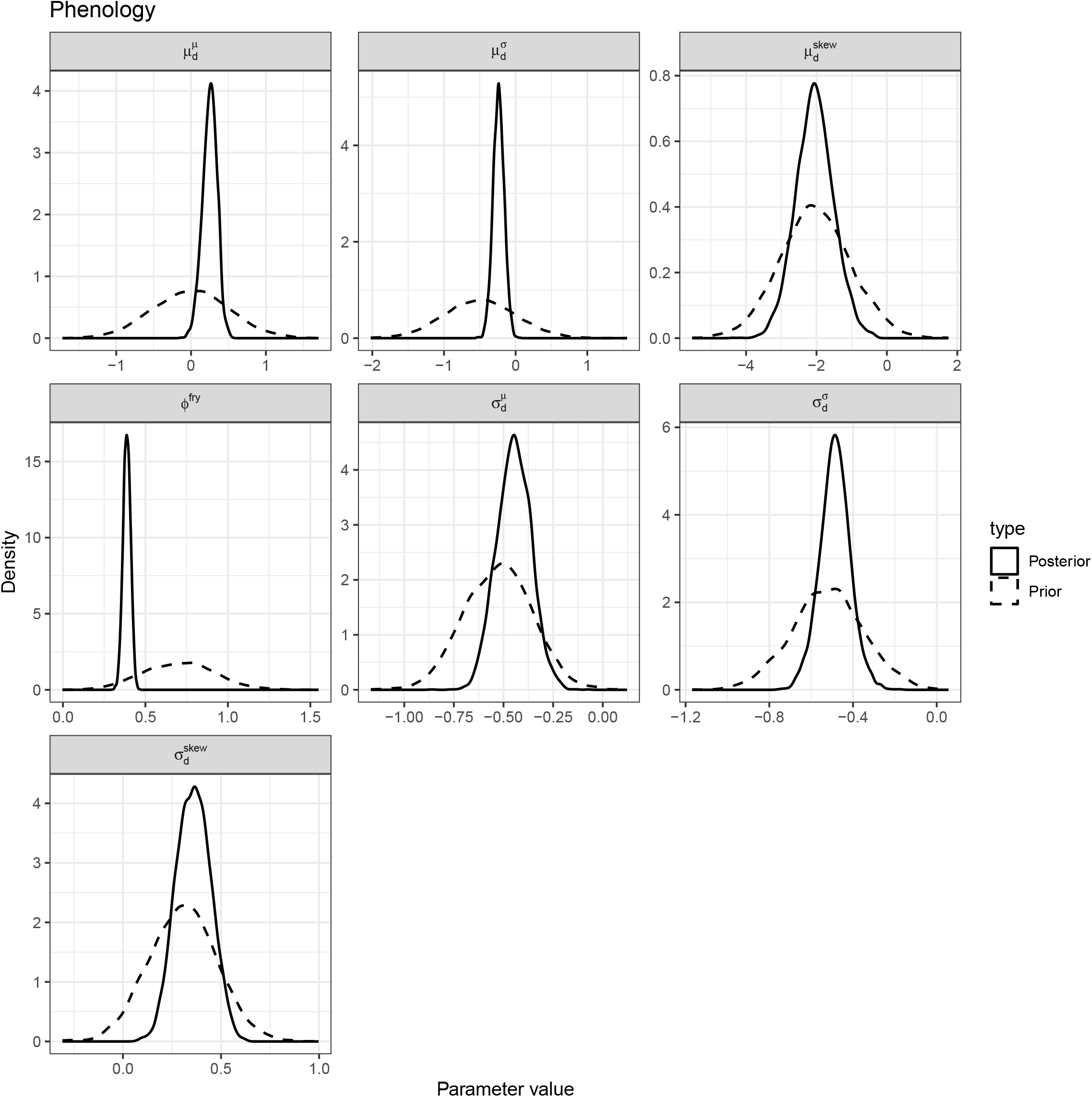
Comparison of prior and posterior for fry emergence phenology parameters.

**Figure S9:**
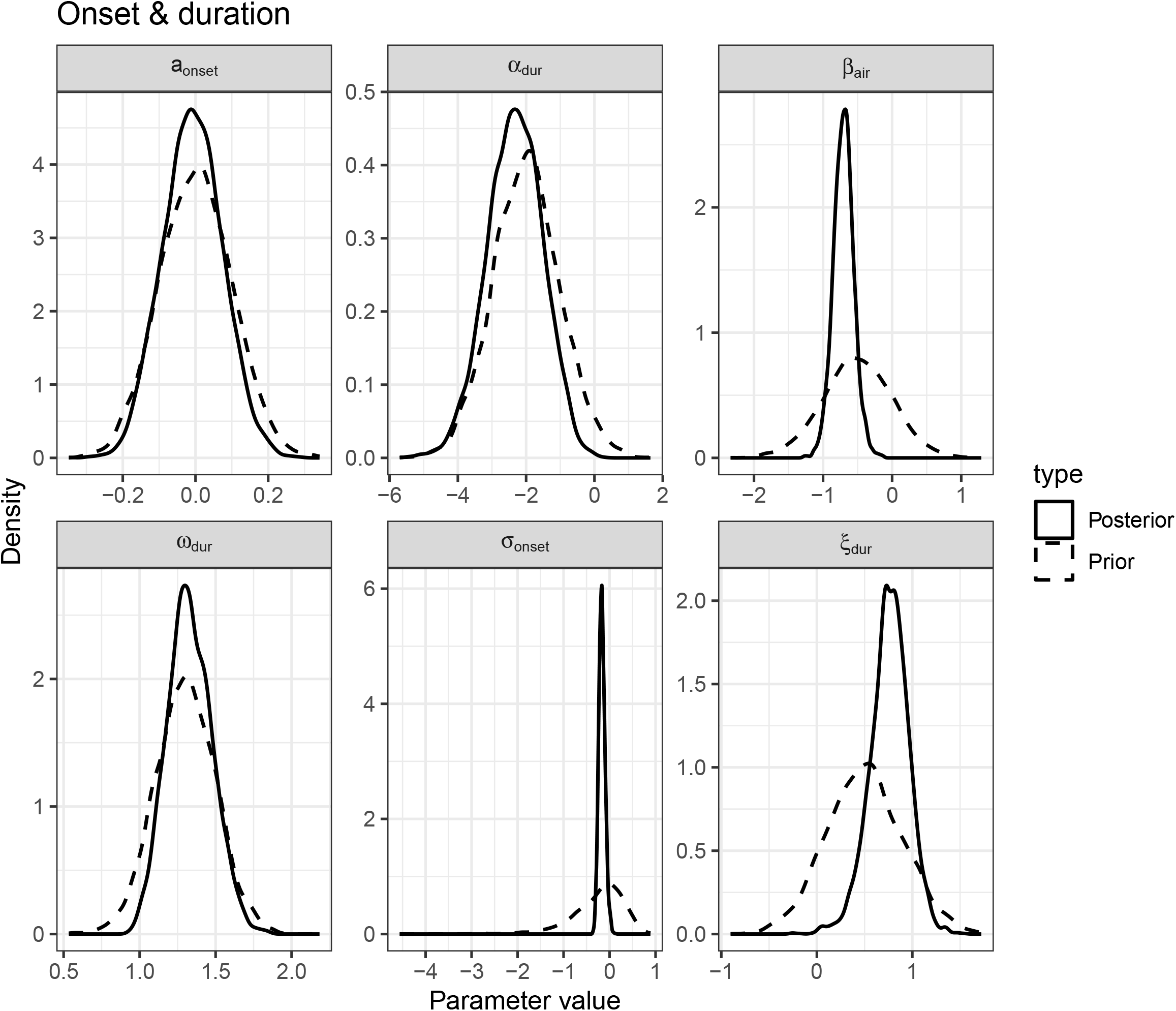
Comparison of prior and posterior for air-water relationship parameters.

## References

Alexander, C.A.D., Alameddine, I., Machin, D., and Alex, K. 2024. A Weight-of-Evidence Approach for Understanding the Recovery of Okanagan Sockeye Salmon. Environmental Management 74(6): 1063–1085. Springer Science and Business Media LLC. doi:10.1007/s00267-024-02031-y.

Bailey, C.J., Stiff, H.W., Thompson, P.L., Judson, B., and Ogden, A.D. 2025. Osoyoos Lake Sockeye Salmon (Oncorhynchus Nerka) return abundance (1977-2023), adult age composition (1985-2023), and marine survival (1998-2021). Can. Manuscr. Rep. Fish. Aquat. Sci. 3282: vi+49 p.

Beacham, T.D., and Murray, C.B. 1990. Temperature, Egg Size, and Development of Embryos and Alevins of Five Species of Pacific Salmon: A Comparative Analysis. Transactions of the American Fisheries Society 119(6): 927–945. Oxford University Press (OUP). doi:10.1577/1548-8659(1990)119<0927:tesado>2.3.co;2.

Beverton, R.J.H., and Holt, S.J. 1957. On the Dynamics of Exploited Fish Populations. Ministry of Agriculture, Fisheries and Food.

Braun, D.C., Patterson, D.A., and Reynolds, J.D. 2013. Maternal and environmental influences on egg size and juvenile life-history traits in Pacific salmon. Ecology and Evolution 3(6): 1727–1740. doi:10.1002/ece3.555.

Brown, N., Stiff, H., Bendriem, N., Luedke, W., Gibeau, P., Bocking, R., Lane, J., McHugh, D., Cunningham, D., Murrell, G., LaFlamme, J., and Dobson, D. 2026. Barkley Sound lake-type Sockeye salmon (Oncorhynchus Nerka) stock assessment in 2025. DFO Can. Sci. Advis. Sec. Res. Doc. 2026(nnn): iv+xx. doi:xxx.xxx.xxx.

Burgner, R.L., DiCostanzo, C.J., Ellis, R.J., Harry, G.Y.Jr., Hartman, W.L., Kerns, O.E.Jr., Mathisen, O.A., and Royce, W.F. 1969. Biological Studies and Estimates of Optimum Escapements of Sockeye Salmon in the Major River Systems in Southwestern Alaska. Alaska. Fish. Bull. U.S. 67: 405–459.

Casas-Mulet, R., Saltveit, S.J., and Alfredsen, K. 2015. The Survival of Atlantic Salmon (SALMO SALAR) Eggs During Dewatering in a River Subjected to Hydropeaking. River Research and Applications 31(4): 433–446. Wiley. doi:10.1002/rra.2827.

Columbia River DART, Columbia Basin Research, University of Washington. 2025. Columbia Basin Adult Passage Historical Run Timing. [accessed 3 October 2025].

Connors, B., Ruggerone, G.T., and Irvine, J.R. 2025. Adapting management of Pacific salmon to a warming and more crowded ocean. ICES Journal of Marine Science 82(1): fsae135. doi:10.1093/icesjms/fsae135.

Crossin, G.T., Hinch, S.G., Cooke, S.J., Welch, D.W., Patterson, D.A., Jones, S.R.M., Lotto, A.G., Leggatt, R.A., Mathes, M.T., Shrimpton, J.M., Van Der Kraak, G., and Farrell, A.P. 2008. Exposure to high temperature influences the behaviour, physiology, and survival of sockeye salmon during spawning migration. Canadian Journal of Zoology 86(2): 127–140. doi:10.1139/Z07-122.

DeCicco, L., Hirsch, R., Lorenz, D., Read, J., Walker, J., Platt, L., Watkins, D., Blodgett, D., Johnson, M., Krall, A., Stanish, L., Zemmels, J., Hinman, E., and Mahoney, M. 2025. dataRetrieval: R packages for discovering and retrieving water data available from U.S. Federal hydrologic web services. Manual, U.S. Geological Survey / U.S. Geological Survey, Reston, VA. doi:10.5066/P9×4L3GE.

DeVries, P. 1997. Riverine salmonid egg burial depths: Review of published data and implications for scour studies. Canadian Journal of Fisheries and Aquatic Sciences 54(8): 1685–1698. Canadian Science Publishing. doi:10.1139/f97-090.

Eliason, E.J., Clark, T.D., Hague, M.J., Hanson, L.M., Gallagher, Z.S., Jeffries, K.M., Gale, M.K., Patterson, D.A., Hinch, S.G., and Farrell, A.P. 2011. Differences in Thermal Tolerance Among Sockeye Salmon Populations. Science 332(6025): 109–112. doi:10.1126/science.1199158.

Essington, T.E., Quinn, T.P., and Ewert, V.E. 2000. Intra- and inter-specific competition and the reproductive success of sympatric Pacific salmon. Canadian Journal of Fisheries and Aquatic Sciences 57(1): 205–213. doi:10.1139/f99-198.

Fenkes, M., Shiels, H.A., Fitzpatrick, J.L., and Nudds, R.L. 2016. The potential impacts of migratory difficulty, including warmer waters and altered flow conditions, on the reproductive success of salmonid fishes. Comparative Biochemistry and Physiology Part A: Molecular & Integrative Physiology 193: 11–21. doi:10.1016/j.cbpa.2015.11.012.

Gabry, J., Češnovar, R., Johnson, A., and Bronder, S. 2024. Cmdstanr: R Interface to ‘CmdStan’. Manual.

Gosselin, J.L., Buhle, E.R., Van Holmes, C., Beer, W.N., Iltis, S., and Anderson, J.J. 2021. Role of carryover effects in conservation of wild Pacific salmon migrating regulated rivers.Ecosphere 12(7): e03618. doi:10.1002/ecs2.3618.

Grant, S.C.H., MacDonald, B.L., and Winston, M.L. 2019. State of Canadian Pacific Salmon: Responses to Changing Climate and Habitats. Can. Tech. Rep. Fish. Aquat. Sci. 3332: ix+50 p.

Hyatt, K.D., Alexander, C.A.D., and Stockwell, M.M. 2015. A decision support system for improving “fish friendly” flow compliance in the regulated Okanagan Lake and River System of British Columbia. Canadian Water Resources Journal / Revue canadienne des ressources hydriques 40(1): 87–110. Informa UK Limited. doi:10.1080/07011784.2014.985510.

Hyatt, K.D., Stiff, H.W., and Stockwell, M.M. 2020. Historic water temperature (1924-2018), river discharge (1929-2018), and adult sockeye salmon migration (1937-2018) observations in the Columbia, Okanogan, and Okanagan rivers. Can. Manuscr. Rep. Fish. Aquat. Sci. 3206: xv + 203 p.

Ingram, S.R. 2011. Ecological implications of flow-mediated scour events for sockeye salmon alevins (Oncorhynchus Nerka). PhD thesis, University of British Columbia.

Lehner, B., and Grill, G. 2013. Global river hydrography and network routing: Baseline data and new approaches to study the world’s large river systems. Hydrological Processes 27(15): 2171–2186. doi:10.1002/hyp.9740.

Martins, E.G., Hinch, S.G., Cooke, S.J., and Patterson, D.A. 2012a. Climate effects on growth, phenology, and survival of sockeye salmon (Oncorhynchus Nerka): A synthesis of the current state of knowledge and future research directions. Reviews in Fish Biology and Fisheries 22(4): 887–914. doi:10.1007/s11160-012-9271-9.

Martins, E.G., Hinch, S.G., Patterson, D.A., Hague, M.J., Cooke, S.J., Miller, K.M., Robichaud, D., English, K.K., and Farrell, A.P. 2012b. High river temperature reduces survival of sockeye salmon (Oncorhynchus Nerka) approaching spawning grounds and exacerbates female mortality. Canadian Journal of Fisheries and Aquatic Sciences 69(2): 330–342. doi:10.1139/f2011-154.

Massicotte, P., and South, A. 2026. Rnaturalearth: World map data from natural earth. Manual.

McElreath, R. 2020. Statistical rethinking: A Bayesian course with examples in R and Stan. In 2nd edition. CRC Press. Taylor and Francis, Boca Raton.

Miller-Saunders, K., Akbarzadeh, A., Fryer, J., and Fleet, J. 2024. Application of genomic tools to validate mechanisms of severe declines in Okanagan Sockeye salmon. Final {{Report}}, Pacific Salmon Commission.

Minke-Martin, V., Hinch, S.G., Braun, D.C., Burnett, N.J., Casselman, M.T., Eliason, E.J., and Middleton, C.T. 2018. Physiological condition and migratory experience affect fitness-related outcomes in adult female sockeye salmon. Ecology of Freshwater Fish 27(1): 296–309. doi:10.1111/eff.12347.

Moring, J.R. 1982. Decrease in stream gravel permeability after clear-cut logging: An indication of intragravel conditions for developing salmonid eggs and alevins. Hydrobiologia 88(3): 295–298. Springer Science and Business Media LLC. doi:10.1007/bf00008510.

Murauskas, J., Hyatt, K., Fryer, J., Koontz, E., Folks, S., Bussanich, R., and Shelby, K. 2021. Migration and survival of Okanagan River Sockeye Salmon Oncorhynchus Nerka, 2012–2019. Animal Biotelemetry 9(1): 37. doi:10.1186/s40317-021-00262-y.

Ng, E., Alex, K., Hyatt, K., Machin, D., Gardner, E., Murauskas, J., and Stockwell, M. 2025. Managing flows for sockeye salmon emergence using the Fish Water Management Tool. Canadian Water Resources Journal/Revue canadienne des ressources hydriques 50(1): 61–78. doi:10.1080/07011784.2025.2459421.

Ogden, A., Alex, K., Pestal, G., Alameddine, I., Davis, B., Judson, B., Stiff, H., and Pham, S. 2025. Wild Salmon Policy Status, Limit Reference Point, and Candidate Escapement Goals for Okanagan Sockeye Salmon. Research {{Document}}, Fisheries and Oceans Canada.

Okanagan Basin Implementation Board. 1982. Report on the Okanagan Basin Implementation Agreement. British Columbia, Canada.

Patterson, D.A., Macdonald, J.S., Hinch, S.G., Healey, M.C., and Farrell, A.P. 2004. The effect of exercise and captivity on energy partitioning, reproductive maturation and fertilization success in adult sockeye salmon. Journal of Fish Biology 64(4): 1039–1059. doi:10.1111/j.1095-8649.2004.0370.x.

Pearl, J., Glymour, M., and Jewell, N.P. 2016. Causal inference in statistics: A primer. Wiley, Chichester, West Sussex.

Quinn, T.P., Eggers, D.M., Clark, J.H., and Rich, H.B., Jr. 2007. Density, climate, and the processes of prespawning mortality and egg retention in Pacific salmon (Oncorhynchus spp.). Canadian Journal of Fisheries and Aquatic Sciences 64(3): 574–582. doi:10.1139/f07-035.

R Core Team. 2024. R: A language and environment for statistical computing. Manual, R Foundation for Statistical Computing, Vienna, Austria.

Rand, P.S., Goslin, M., Gross, M.R., Irvine, J.R., Augerot, X., McHugh, P.A., and Bugaev, V.F. 2012. Global Assessment of Extinction Risk to Populations of Sockeye Salmon Oncorhynchus Nerka. PLoS ONE 7(4): e34065. Public Library of Science (PLoS). doi:10.1371/journal.pone.0034065.

Rand, P.S., Hinch, S.G., Morrison, J., Foreman, M.G.G., MacNutt, M.J., Macdonald, J.S., Healey, M.C., Farrell, A.P., and Higgs, D.A. 2006. Effects of River Discharge, Temperature, and Future Climates on Energetics and Mortality of Adult Migrating Fraser River Sockeye Salmon. Transactions of the American Fisheries Society 135(3): 655–667. doi:10.1577/T05-023.1.

Reiser, D.W., Ramey, M.P., Beck, S., Lambert, T.R., and Geary, R.E. 1989. Flushing flow recommendations for maintenance of salmonid spawning gravels in a steep, regulated stream. Regulated Rivers: Research & Management 3(1): 267–275. Wiley. doi:10.1002/rrr.3450030126.

Ricker, W.E. 1954. Stock and Recruitment. Journal of the Fisheries Research Board of Canada 11(5): 559–623. doi:10.1139/f54-039.

Shepherd, J.G. 1982. A versatile new stock-recruitment relationship for fisheries, and the construction of sustainable yield curves. ICES Journal of Marine Science 40(1): 67–75. doi:10.1093/icesjms/40.1.67.

Sopinka, N.M., Middleton, C.T., Patterson, D.A., and Hinch, S.G. 2016. Does maternal captivity of wild, migratory sockeye salmon influence offspring performance? Hydrobiologia 779(1): 1–10. doi:10.1007/s10750-016-2763-1.

Stan Development Team. 2024. Stan modeling language users guide and reference manual. Manual.

Stiff, H.W., and Thompson, P.L. 2026. Projected water temperature and exposure indices for adult sockeye salmon migration in the Okanagan watershed, 2010–2099. Canadian Technical Report of Fisheries and Aquatic Sciences xxxx: yyy–zzz. Fisheries and Oceans Canada.

Stockwell, M.M., Hyatt, K.D., Alex, K., Louie, C., and Machin, D. 2020. Methods and Summary Observations of Okanagan Sockeye Salmon Spawn Timing, Fry Emergence, and Associated Water Temperatures (Brood Years 2002-2018). Can. Data. Rep. Fish. Aquat. Sci. 1300: vii + 61 p.

Summit Environmental Consultants Ltd. 2002. Observations & modelling of bed movement & redd scour in Okanagan River. Vernon, Canada.

Thompson, P.L., Akenhead, S.A., and Louie, C. In press. Estimating spawning salmon abundance from visual surveys: A hierarchical model of arrival and exit dynamics. in review.

Thorstad, E.B., Bliss, D., Breau, C., Damon-Randall, K., Sundt-Hansen, L.E., Hatfield, E.M.C., Horsburgh, G., Hansen, H., Maoiléidigh, N.Ó., Sheehan, T., and Sutton, S.G. 2021. Atlantic salmon in a rapidly changing environment—Facing the challenges of reduced marine survival and climate change. Aquatic Conservation: Marine and Freshwater Ecosystems 31(9): 2654–2665. doi:10.1002/aqc.3624.

Vehtari, A., Gelman, A., Simpson, D., Carpenter, B., and Bürkner, P.-C. 2021. Rank-Normalization, Folding, and Localization: An Improved R^ for Assessing Convergence of MCMC (with Discussion). Bayesian Analysis 16(2). doi:10.1214/20-BA1221.

Whitney, C.K., Hinch, S.G., and Patterson, D.A. 2013. Provenance matters: Thermal reaction norms for embryo survival among sockeye salmon Oncorhynchus Nerka populations. Journal of Fish Biology 82(4): 1159–1176. doi:10.1111/jfb.12055.

Wilson, S.M., and Peacock, S.J. 2025. Freshwater life-cycle timing of Pacific salmon and steelhead (Oncorhynchus spp.) in Canada. Canadian Journal of Fisheries and Aquatic Sciences 82: 1–17. Canadian Science Publishing. doi:10.1139/cjfas-2024-0213.

